# Human organ chip-enabled pipeline to rapidly repurpose therapeutics during viral pandemics

**DOI:** 10.1101/2020.04.13.039917

**Authors:** Longlong Si, Haiqing Bai, Melissa Rodas, Wuji Cao, Crystal Yuri Oh, Amanda Jiang, Rasmus Moller, Daisy Hoagland, Kohei Oishi, Shu Horiuchi, Skyler Uhl, Daniel Blanco-Melo, Randy A. Albrecht, Wen-Chun Liu, Tristan Jordan, Benjamin E. Nilsson-Payant, James Logue, Robert Haupt, Marisa McGrath, Stuart Weston, Atiq Nurani, Seong Min Kim, Danni Y. Zhu, Kambez H. Benam, Girija Goyal, Sarah E. Gilpin, Rachelle Prantil-Baun, Rani K. Powers, Kenneth Carlson, Matthew Frieman, Benjamin R. tenOever, Donald E. Ingber

## Abstract

The rising threat of pandemic viruses, such as SARS-CoV-2, requires development of new preclinical discovery platforms that can more rapidly identify therapeutics that are active *in vitro* and also translate *in vivo*. Here we show that human organ-on-a-chip (Organ Chip) microfluidic culture devices lined by highly differentiated human primary lung airway epithelium and endothelium can be used to model virus entry, replication, strain-dependent virulence, host cytokine production, and recruitment of circulating immune cells in response to infection by respiratory viruses with great pandemic potential. We provide a first demonstration of drug repurposing by using oseltamivir in influenza A virus-infected organ chip cultures and show that co-administration of the approved anticoagulant drug, nafamostat, can double oseltamivir’s therapeutic time window. With the emergence of the COVID-19 pandemic, the Airway Chips were used to assess the inhibitory activities of approved drugs that showed inhibition in traditional cell culture assays only to find that most failed when tested in the Organ Chip platform. When administered in human Airway Chips under flow at a clinically relevant dose, one drug – amodiaquine - significantly inhibited infection by a pseudotyped SARS-CoV-2 virus. Proof of concept was provided by showing that amodiaquine and its active metabolite (desethylamodiaquine) also significantly reduce viral load in both direct infection and animal-to-animal transmission models of native SARS-CoV-2 infection in hamsters. These data highlight the value of Organ Chip technology as a more stringent and physiologically relevant platform for drug repurposing, and suggest that amodiaquine should be considered for future clinical testing.

---

The increasing incidence of potential pandemic viruses, such as influenza A virus, Middle East respiratory syndrome coronavirus (MERS-CoV), Severe acute respiratory virus (SARS-CoV), and now SARS-CoV-2, requires development of new preclinical approaches that can accelerate development of effective therapeutics and prophylactics. The most rapid way to confront a pandemic challenge would be to repurpose existing drugs that are approved for other medical indications as antiviral therapeutics. While researchers and clinicians around the world are attempting to do this for the COVID-19 pandemic, current approaches have been haphazard and generally rely entirely on results of *in vitro* screens with cell lines. This has resulted in equivocal results regarding drug efficacies and possible toxicity risks as in the case of hydroxychloroquine and chloroquine^1–4^, and thus, there is a great need to address this problem in a more systematic and human-relevant way. Recognizing the potential danger of unforeseen pandemics over two years ago, the Defense Advanced Research Projects Agency (DARPA) and National Institutes of Health (NIH) funded work to explore whether human organ-on-a-chip (Organ Chip) microfluidic culture technology might be helpful in confronting potential biothreat challenges. We previously showed that Organ Chips can recapitulate human organ physiology, disease states, and therapeutic responses to clinically relevant drug exposures with high fidelity^5–9^. Here we show how human Lung Airway Chips may be used to model human lung responses to viral infection *in vitro*, and in concert with higher throughput cell-based assays and animal models, to identify existing approved drugs that have the potential to be repurposed for treating or preventing spread of viral pandemics caused by influenza A virus or SARS-CoV-2.

Infections by respiratory viruses and antiviral drug screening assays are currently studied *in vitro* using cultured established cell lines, primary tissue-derived human cells, human organoids, and *ex vivo* human lung tissue cultures despite all having significant limitations (**Supplementary Table 1**)^10–13^. For example, established cell lines commonly demonstrate various defects in their capacity to elicit an antiviral state because normally elicited interferons cause cell cycle arrest, and so this dynamic is often selected against with continuous passaging^14^. Arguably even more important, cell lines and even human primary cells grown in conventional cultures do not exhibit the highly differentiated tissue structures and functions (e.g., mucociliary clearance) seen in living human organs. Explant cultures of human respiratory tract tissue circumvent this limitation, but their availability is limited and their viability can only be maintained for a short time (4-10 days)^12, 15^. While human lung organoids provide a more functional lung epithelium, they do not allow culturing of the epithelium at an air-liquid interface (ALI) or modeling of other physiologically relevant organ-level features of lung, such as mucus layer formation, mucociliary clearance, cross-talk between epithelium and endothelium, or recruitment of circulating immune cells^10, 11^, all of which play key roles in host responses to infection by respiratory viruses. Moreover, in all of these culture models, drug studies are carried out under static conditions that cannot predict human responses to clinically relevant, dynamic drug exposure profiles that result from complex pharmacokinetics (PK) *in vivo*. Thus, there is an urgent need for alternative preclinical *in vitro* models that specifically mimic human lung responses to infection by potential pandemic respiratory viruses, and because of their ability to recapitulate human organ-level physiology and pathophysiology, human Lung Chips offer a potential solution.

## Influenza A virus infection and immune responses replicated in human Lung Chips

To initially assess whether Organ Chip technology^5–8^ can be used to create a preclinical *in vitro* model for therapeutics discovery, we tested it against a drug that is used clinically for treatment of influenza A viral infections. The human Lung Airway Chip is a microfluidic device that contains two parallel microchannels separated by an extracellular matrix (ECM)-coated porous membrane (**Fig. 1a**)^16^. Primary human lung airway basal stem cells are grown under an air-liquid interface (ALI) on one side of the membrane in the ‘airway channel’, while interfaced with a primary human lung endothelium grown on the opposite side of the same membrane, which is exposed to continuous fluid flow of culture medium within the parallel ‘vascular channel’ (**Fig. 1a**). This device supports differentiation of the lung airway basal stem cells into a mucociliary, pseudostratified epithelium with proportions of airway-specific cell types (ciliated cells, mucus-producing goblet cells, club cells, and basal cells) (**Extended Data Fig. 1a**), as well as establishment of continuous ZO1-containing tight junctions and cilia (**Fig. 1b**), permeability barrier properties, and mucus production (**Extended Data Fig. 1b,c**) similar to those observed in human airway *in vivo*^17^, as well as in prior Airway Chip studies that used a membrane with smaller pores that did not permit immune cell transmigration^16^. The underlying human pulmonary microvascular endothelium also forms a continuous planar cell monolayer with cells linked by VE-cadherin containing adherens junctions (**Fig. 1b**) as it does *in vivo*.

**Fig. 1.**
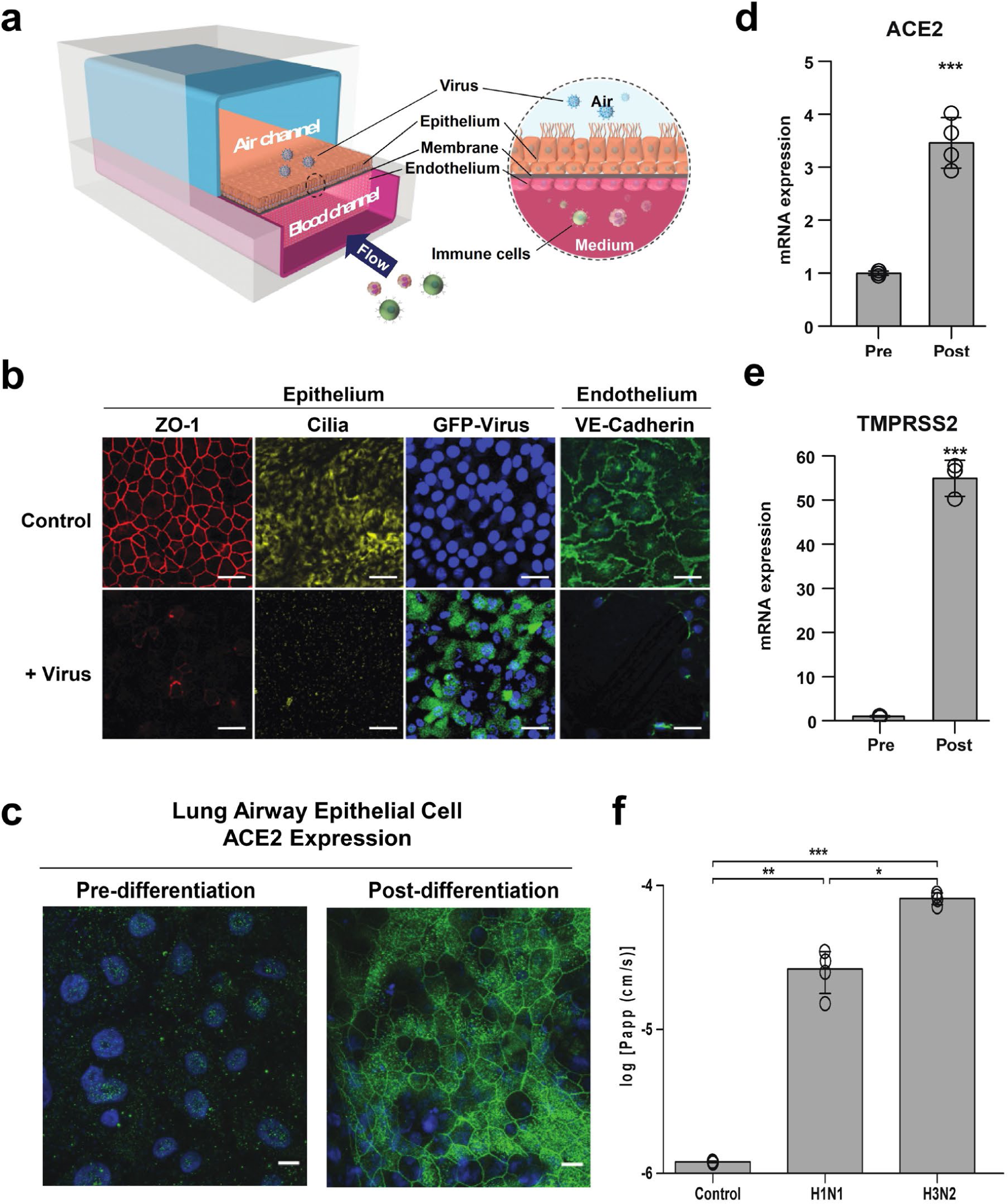
Characterization of the human Airway Chip and its infection with influenza virus. (**a**) Schematic diagram of a cross-section through the Airway Chip. (**b**) Immunofluorescence micrographs showing the distribution of ZO1-containing tight junctions and cilia in the epithelium and VE-cadherin-containing adherens junctions in the endothelium of the Airway Chip in the absence (Control) or presence (+ Virus) of infection with GFP-labeled influenza PR8 (H1N1) virus (MOI = 0.1) for 48 h (blue, DAPI-stained nuclei; bar, 50 μm). (**c-e**) Immunofluorescence micrographs showing the expression of ACE2 receptor (**c**) and fold changes in mRNA levels of ACE2 (**d**) and TMPRSS2 (**e**) in the well-differentiated primary human lung airway epithelim on-chip (Post) versus the same cells prior to differentiation (Pre). (**f**) Increase in barrier permeability as measured by apparent permeability (log *P_app_*) within the human Airway chip 48 h post-infection with PR8 (H1N1) or HK/68 (H3N2) virus (MOI = 0.1) compared to no infection (Control).

Importantly, the highly differentiated airway epithelium in the Airway Chip expresses higher levels of expression of genes encoding multiple serine proteases involved in viral entry including TMPRSS2, TMPRSS4, TMPRSS11D, and TMPRSS11E (DESC1) compared to MDCK cells that are often used to study influenza virus infection *in vitro* (**Extended Data Fig. 1d**); these proteases are essential for the activation and propagation of influenza viruses *in vivo*. In addition, compared to their initial state upon seeding, differentiation of the airway epithelial cells at an ALI on-chip is accompanied by large increases in protein (**Fig. 1c**) and mRNA expression levels of the SARS-CoV-2 receptor, angiotensin converting enzyme-2 (ACE-2) (**Fig. 1d**) and the TMPRSS2 protease (**Fig. 1e**) that mediate infection by SARS-CoV-2^18, 19^.

When GFP-labeled influenza A/PuertoRico8/34 (H1N1) virus was introduced into the air channel of the microfluidic chip to mimic *in vivo* infection with airborne virus (**Fig. 1a**), real-time fluorescence microscopic analysis confirmed that the virus infected the human airway epithelial cells (**Fig. 1b, Supplementary Movie 1**), and this was accompanied by damage to the epithelium, including disruption of tight junctions, loss of apical cilia (**Fig. 1b)**, and compromised barrier function (**Fig. 1f**). Significantly less infection was detected in undifferentiated airway basal epithelium prior to culture at an ALI on-chip, and there was no detectable direct infection of the endothelium by the virus (**Extended Data Fig. 2a**). Interestingly, however, influenza A virus infection led to disruption of the lung endothelium on-chip, as evidenced by loss of VE-cadherin containing adherens junctions (**Fig. 1b**), which is consistent with the vascular leakage that is induced in lungs of human patients with influenza^20^.

**Fig. 2.**
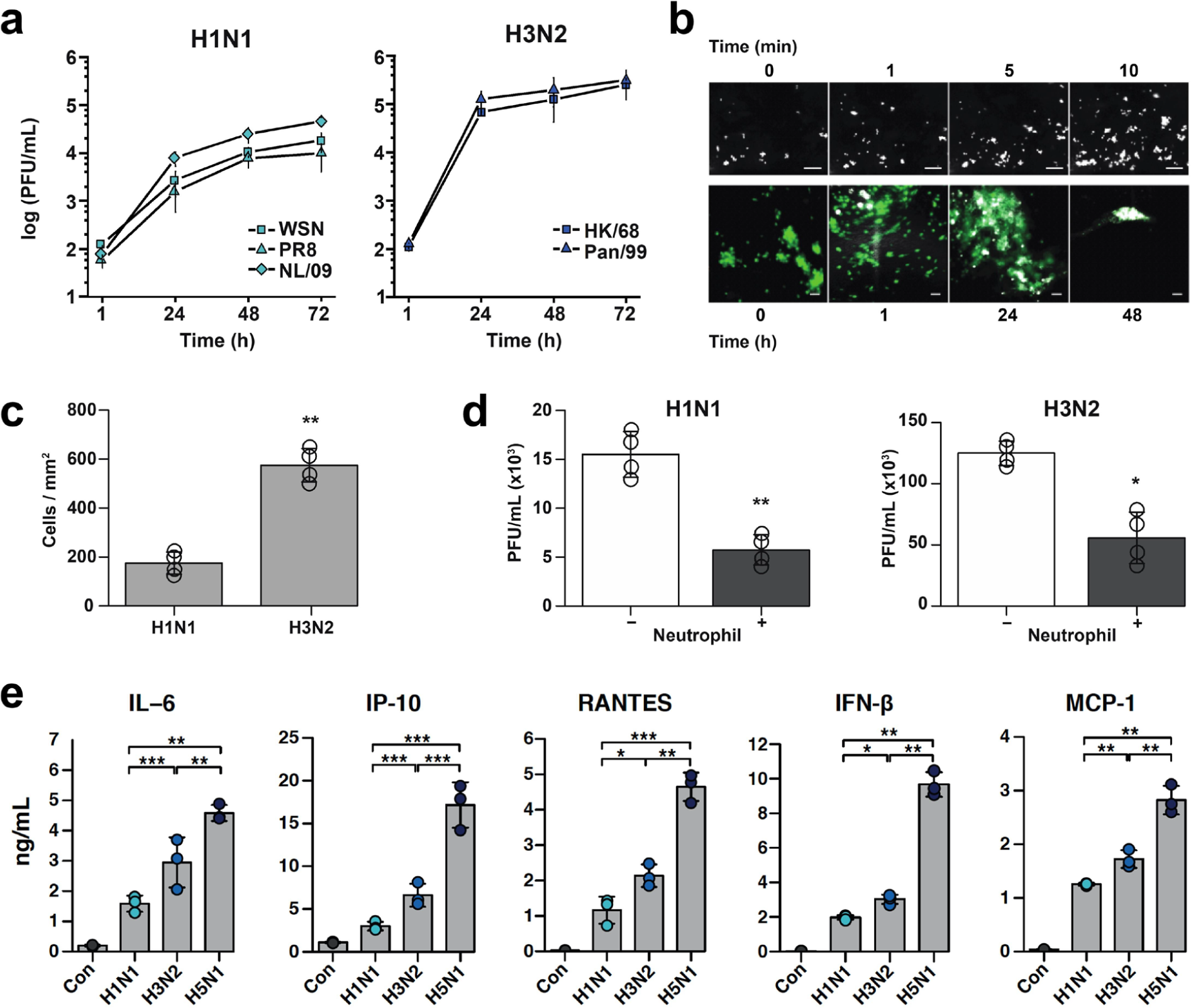
Infection with multiple influenza strains in human Airway Chips and resultant immune responses. (**a**) Replication kinetics of influenza H1N1 virus strains WSN (square), PR8 (triangle), NL/09 (diamond) (left graph), and of influenza H3N2 virus strains HK/68 (square) and Pan/99 (triangle) (right graph), when infected at MOI = 0.001 in human Airway Chips. (**b**) Neutrophil responses to influenza infection in human lung Airway Chip. Top, sequential immunofluorescence micrographs showing time-dependent recruitment of neutrophils (white) to the apical surface of the endothelium (unlabeled) within a human Airway Chip infected with influenza PR8 (H1N1) virus (bar, 50 µm). Bottom, immunofluorescence micrographs showing time-dependent recruitment of neutrophils (white) to the epithelium (unlabeled) and clearance of clustered epithelial cells infected with GFP-labeled PR8 (H1N1) virus (green) (bar, 50 µm). (**c**) Graph showing numbers of neutrophils recruited to the epithelium in response to infection by H1N1 or H3N2. (**d**) Virus titers of human Airway Chips infected with WSN (H1N1) or HK/68 (H3N2) in the presence (+) or absence (-) of added neutrophils (PFU, plaque-forming units). (**e**) Production of indicated cytokines and chemokines in the human Airway chip at 48 h post-infection with different clinically isolated influenza virus strains, including NL/09 (H1N1), Pan/99 (H3N2), and HK/97 (H5N1) (MOI = 0.1). *, P<0.05; **, P<0.01; ***, P<0.001.

Analysis of the replication kinetics of five different influenza A virus strains, including clinical isolates [A/Netherlands/602/2009 (NL/09; H1N1), A/HongKong/8/68 (HK/68; H3N2), A/Panama/2007/99 (Pan/99; H3N2)] and cell culture strains [influenza Puerto Rico/8/34 (H1N1), A/WSN/1933 (WSN; H1N1)], showed that all of the virus variants propagate efficiently as demonstrated by large (10^3^- to 10^4^-fold) increases in viral titers over 24 to 48 hours in highly differentiated human lung airway epithelium on-chip (**Fig. 2a**). Notably, the H3N2 virus strains (HK/68 and Pan/99) exhibited ∼10-fold greater replication efficiency than the H1N1 strains (PR8, WSN, and NL/09) (**Fig. 2a**) and caused severe barrier disruption (**Fig. 1f**) and more cilia loss (**Extended Data Fig. 2b**). These results corroborate the finding that H3N2 is more infectious and virulent, and causes more severe clinical symptoms in humans^21^. Donor-to-donor variability in terms of sensitivity to influenza virus infection was minimal in these studies, as similar viral infectivity was obtained in chips derived from five different healthy epithelial cell donors (**Extended Data Fig. 2c**).

Recruitment of circulating immune cells, such as neutrophils, under dynamic flow to the site of infection in the airway epithelium contributes significantly to influenza A virus pathogenesis in the lung^22^; however, this process has not been well investigated in existing *in-vitro* models due to their 2D static nature. When primary human neutrophils were perfused through the vascular channel of Airway Chips infected with H1N1 or H3N2 virus, we observed recruitment of these circulating immune cells to the apical surface of the activated lung endothelium within minutes (**Fig. 2b top, Supplementary Movie 2**). This was followed by transmigration of the neutrophils through the endothelium and the ECM-coated pores of the intervening membrane, and up into the airway epithelium over hours (**Fig. 2b bottom**). The neutrophils targeted the influenza A virus nucleoprotein (NP)-positive infected airway cells (**Extended Data Fig. 3**) and induced them to coalesce into clusters that decreased in size over time, resulting in clearance of the virus, as evidenced by the disappearance of GFP-positive cells over a period of 1-2 days (**Fig. 2b bottom**). Consistent with the ability of H3N2 virus to induce stronger inflammation relative to H1N1 *in vivo*^21^, H3N2 also stimulated more neutrophil recruitment than H1N1 (**Fig. 2c**), and neutrophil infiltration into the epithelium significantly decreased the viral titers of both H1N1 and H3N2 on-chip (**Fig. 2d**), consistent with the protective role that neutrophils provide by clearing virus *in vivo*^22^. H1N1 infection also was accompanied by increased secretion of various inflammatory cytokines and chemokines, including IL-6, IP-10, RANTES, interferon-β, and MCP-1, which could easily be measured in the effluent from the vascular channel (**Fig. 2e**).

**Fig. 3.**
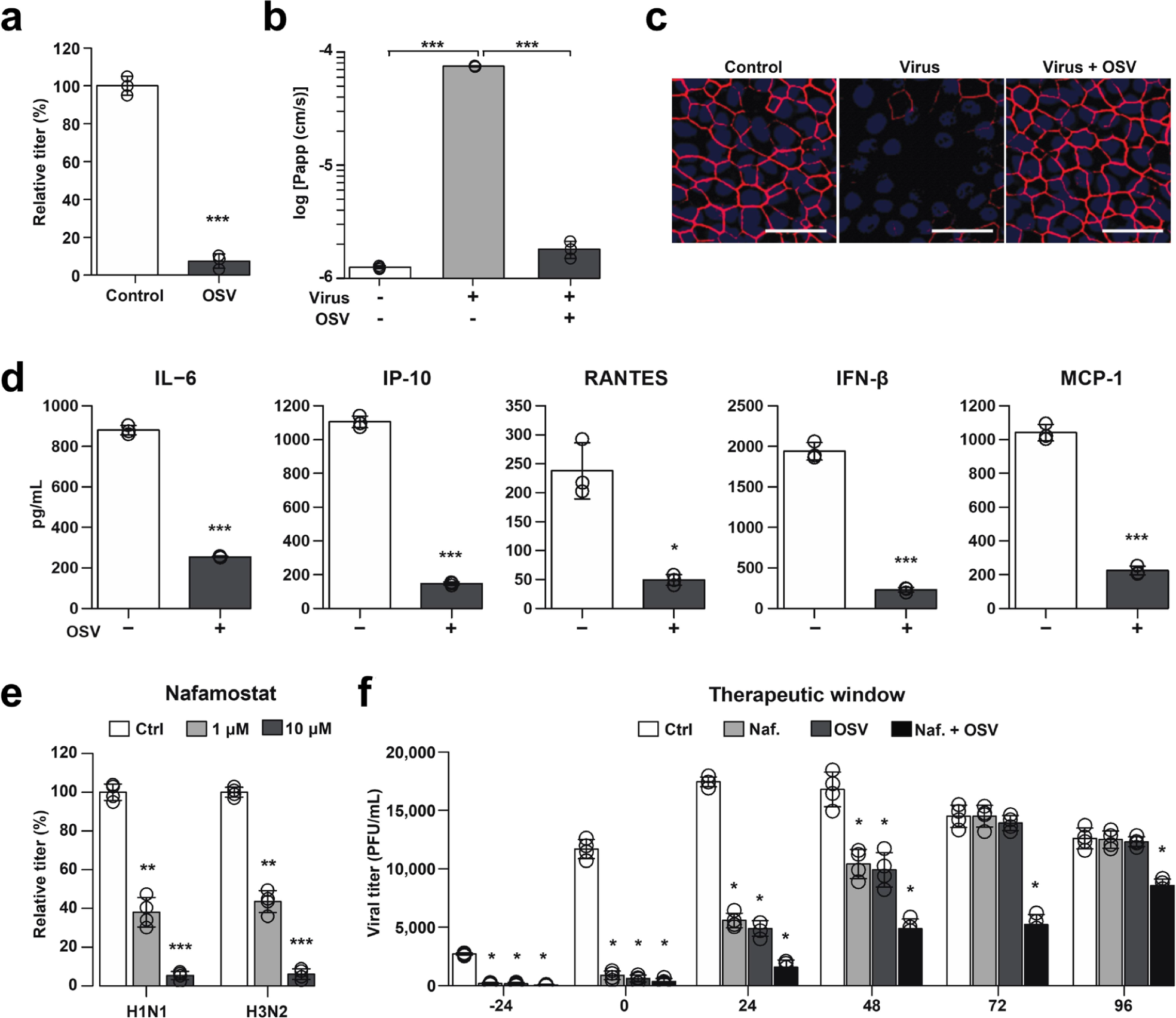
Effects of anti-influenza therapeutics in the human Airway Chip. (**a**) Graph showing relative plaque titers of progeny virus in the absence (Ctrl) or presence of 1 µM oseltamivir acid (OSV) 48 h post-infection with WSN (H1N1; MOI = 0.1). (**b**) Barrier permeability (log P_app_) measured under control conditions within the human Airway Chip (− Virus) or 48 h post-infection with WSN (+ Virus) with (+) or without (-) OSV. (**c**) Immunofluorescence micrographs showing the distribution of ZO1-containing tight junctions in airway epithelium under baseline conditions (Ctrl) or infected with WSN alone (Virus) or in the presence of OSV (Virus + OSV) 48 h post-infection (bar, 50 µm). (**d**) Production of cytokines in human Airway Chip 48 h post-infection with WSN in the presence (+) or absence (-) of OSV. (**e**) Virus titer detection showing the effects of Nafamostat at 1 µM (grey bars) or 10 μM (white bars) dose on virus replication of H1N1 and H3N2 in Airway chips 48 h post-infection compared to untreated chips (Ctrl, black bars). (**f**) The effects of Nafamostat, oseltamivir and their combination on relative viral titers when added to H1N1 virus-infected human Airway Chips at indicated times; note the synergistic effects of these two drugs at later times. *, P<0.05; **, P<0.01; ***, P<0.001.

Variations in secretion of proinflammatory mediators in the human lung airway contribute to differences in pathogenesis and morbidity observed for different influenza A virus strains, and analysis of cytokine levels can help clinicians assess disease severity. Thus, we compared the innate immune responses of the human Airway Chip to infection with three patient-derived influenza A virus strains with different virulence: NL/09 (H1N1), Pan/99 (H3N2), and A/HongKong/156/1997 (HK/97; H5N1). When chips were infected with H3N2 and H5N1 viruses that are known to produce more severe clinical symptoms than H1N1 in patients, we found that they also stimulated production of higher levels of cytokines and chemokines, and the most virulent H5N1 strain induced the highest concentrations (**Fig. 2e**). These results mirror the clinical finding that patients infected with H5N1 have increased serum concentrations of these inflammatory factors relative to those with H1N1 or H3N2, which significantly contributes to disease pathogenesis^21^.

## Recapitulation of the effects of clinically used anti-viral therapeutics

To explore whether the Airway Chip can be used to evaluate the efficacy of potential antiviral therapeutics, we first tested oseltamivir (Tamiflu), which is the anti-influenza drug most widely used in the clinic. As oseltamivir is metabolized by the liver to release oseltamivir acid *in vivo*, we introduced this active metabolite into the vascular channel of Airway Chip infected with H1N1 virus, mimicking its blood levels after oral administration. Oseltamivir (1 µM) efficiently inhibited influenza A virus replication (**Fig. 3a**), prevented virus-induced compromise of barrier function (**Fig. 3b**) and disruption of epithelial tight junctions (**Fig. 3c**), and decreased production of multiple cytokines and chemokines on-chip (**Fig. 3d**). Importantly, similar anti-influenza efficacy was detected in a randomized controlled trial where treatment with Oseltamivir also led to one log drop in viral titers in nasopharyngeal samples provided by 350 patients^23^. Thus, the Airway Chip faithfully replicates the effects of oseltamivir previously observed in humans, suggesting that it may serve as a useful preclinical model to evaluate potential therapies for virus-induced human lung infections in a preclinical setting.

## Repurposing of approved drugs as potential anti-influenza therapeutics

Given that host serine proteases on human airway epithelial cells play critical roles in influenza A virus propagation^12, 24^, and their expression is significantly elevated in the differentiated Airway Chip (**Fig. 1e, Extended Data Fig. 1d**), we explored whether existing approved drugs that inhibit serine proteases could suppress infection by delivering them into the airway channel of influenza virus-infected chips (e.g, to mimic intratracheal delivery by aerosol, nebulizer, or inhaler). These studies revealed that two clinically used anticoagulant drugs, nafamostat (**Fig. 3e**) and trasylol (**Extended Data Fig. 4a**), significantly reduced influenza H1N1 and H3N2 titers on-chip. Further exploration of nafamostat’s actions revealed that it protects airway barrier function (**Extended Data Fig. 4b**) and tight junction integrity (**Extended Data Fig. 4c**), and decreases production of cytokines and chemokines (**Extended Data Fig. 4d**). Nafamostat and the other protease inhibitors appeared to act by efficiently blocking the serine proteases-TMPRSS11D and TMPRSS2-mediated enzymatic cleavage of influenza A viral HA0 protein into HA1 and HA2 subunits (**Extended Data Fig. 4e**), which is required for viral entry^25^.

**Fig. 4.**
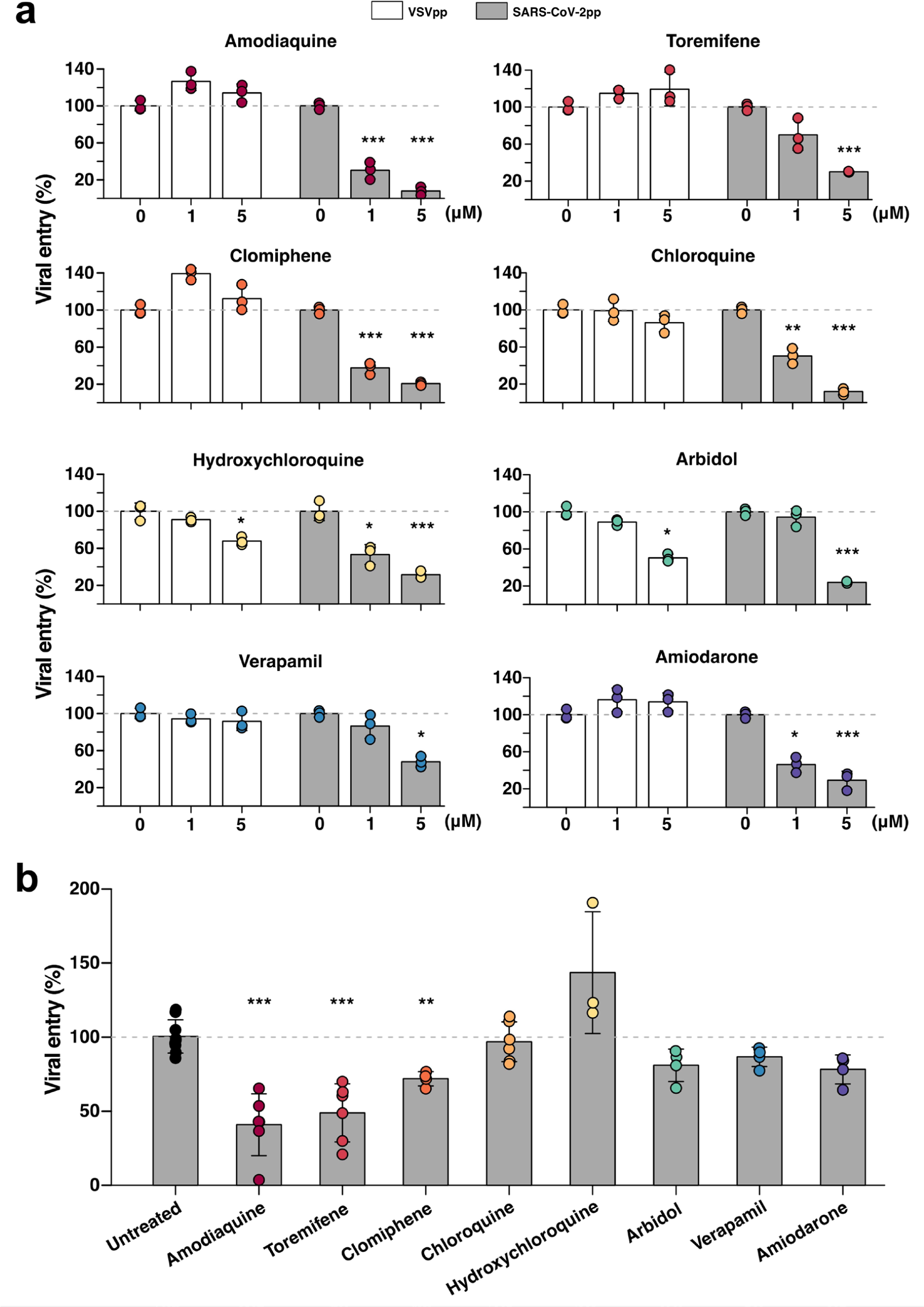
Effects of FDA-approved drugs on pseudotyped SARS-CoV-2 viral entry in Huh-7 cells versus human Airway Chips. (**a**) Graphs showing the inhibitory effects of amodiaquine, toremifene, clomiphene, chloroquine, hydroxychloroquine, arbidol, verapamil, and amiodarone when added at 0, 1, or 5 µM to Huh-7 cells infected with SARS-CoV-2pp for 72 h (grey bars). The number of pseudoparticles in the infected cells was quantified by measuring luciferase activity; viral entry in untreated cells was set as 100%. VSVpp were tested in parallel to exclude toxic and nonspecific effects of the drugs tested (white bars). (**b**) The efficacy of the same drugs in human Airway Chips infected with SARS-CoV-2pp. Amodiaquine, toremifene, clomiphene, chloroquine, hydroxychloroquine, arbidol, verapamil, and amiodarone were delivered into apical and basal channels of the chip at their respective C_max_ in human blood, and one day later chips were infected with SARS-CoV-2pp while in the continued presence of the drugs for 2 more days. The epithelium from the chips were collected for detection of viral pol gene by qRT-PCR; viral entry in untreated chips was set as 100%. *, P < 0.05; **, P < 0.01; ***, P < 0.001.

When we added nafamostat or oseltamivir at different time points during influenza virus infection on-chip, both nafamostat and oseltamivir exhibited prophylactic and therapeutic effects (**Fig. 3f**). However, oseltamivir only produced therapeutic effects when was administered within 48 h post-infection (**Fig. 3f**). This is consistent with the observation that oseltamivir is only recommended for clinical use within 2 days of influenza virus infection^26^, which is one of the important limitations of using this antiviral therapeutic clinically. Nafamostat also exhibited its inhibitory effects over a 48 h time period (**Fig. 3f**). Impressively, however, combined administration of nafamostat and oseltamivir exerted more potent inhibition of influenza virus infection, and this combined regimen was able to double oseltamivir’s treatment time window from 48 to 96 hours (**Fig. 3f**).

## Identification of approved drugs as SARS-CoV-2 entry inhibitors

Given the faithful recapitulation of human lung responses to influenza infection, we quickly pivoted our effort to focus on SARS-CoV-2 infection when we learned of the emerging COVID-19 pandemic. To alleviate safety concerns and immediately initiate work in a BSL2 laboratory, we designed SARS-CoV-2 pseudoparticles (SARS-CoV-2pp) that contain the SARS-CoV-2 spike (S) protein assembled onto luciferase reporter gene-carrying retroviral core particles^27^, based on the genome sequence of SARS-CoV-2 released in GenBank on January 12, 2020^28^. We confirmed the incorporation of SARS-CoV-2 S protein into SARS-CoV-2pp by Western blotting (**Extended Data Fig. 5a**), as previously shown in other pseudotyped SARS-CoV-2 viruses^18^. Successful generation of SARS-CoV-2pp was further confirmed by efficient infection in Huh-7 cells, a human liver cell line commonly used to study infection of SARS viruses^29^, whereas control pseudoparticles without the spike protein of SARS-CoV-2 did not infect **(Extended Data Fig. 5b**). These pseudotyped S protein-expressing viral particles faithfully reflect key aspects of native SARS-CoV-2 entry into host cells via binding to its ACE2 receptor^30^, and thus, they can be used to test potential entry inhibitors of SARS-CoV-2^18, 27^. Vesicular stomatitis virus (VSV) GP protein pseudoparticles (VSVpp) were also generated and used in parallel studies to exclude toxic and non-specific effects of SARS-CoV-2 entry inhibitors^18, 27^.

**Fig. 5.**
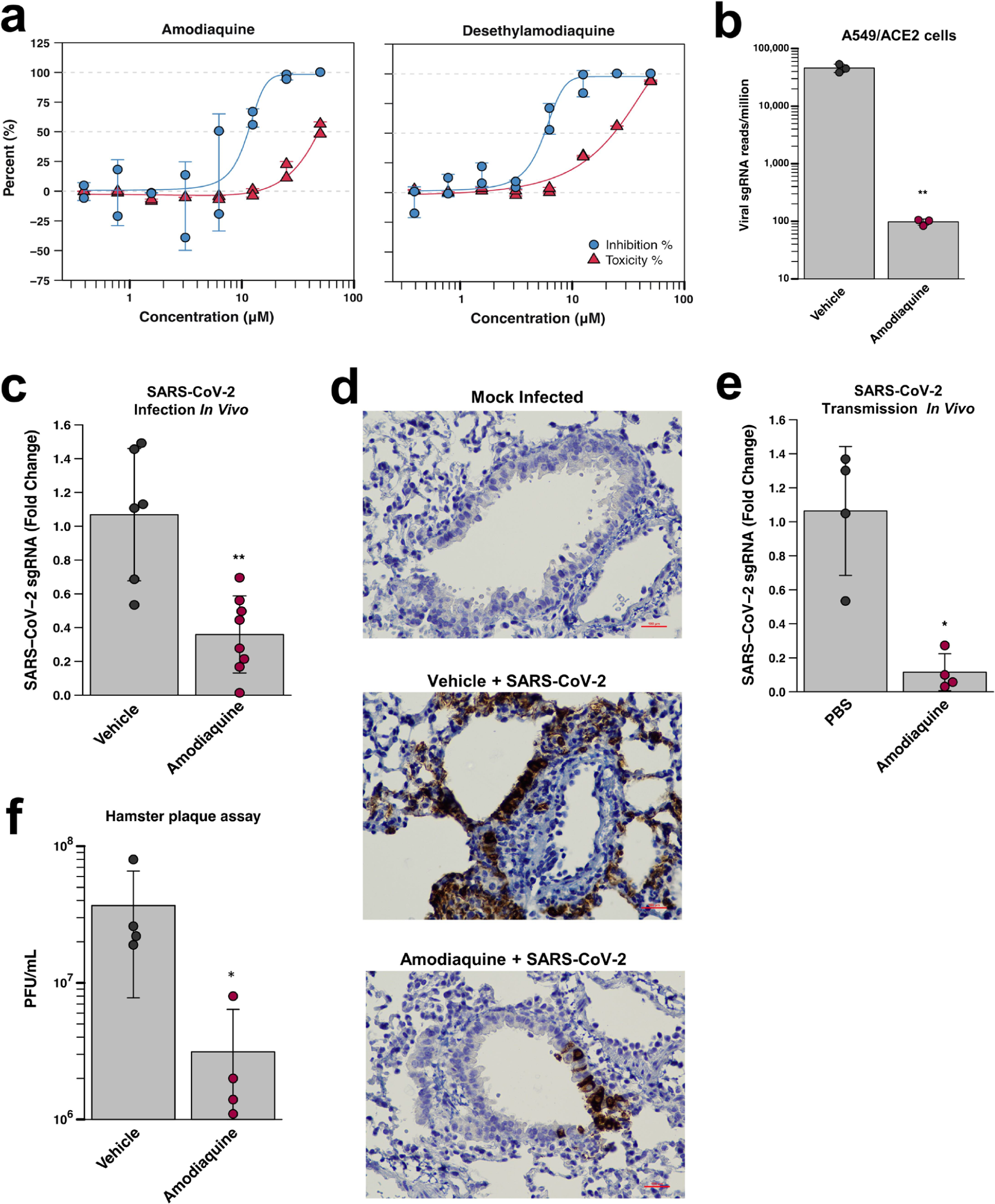
Inhibition of infection by native SARS-CoV-2 virus *in vitro* and *in vivo.* (**a**) Dose-response curves for amodiaquine and its metabolite desethylamodiaquine demonstrating their ability to inhibit GFP-SARS-CoV-2 infection (MOI = 0.1) in a dose-dependent manner in Vero E6 cells. (**b**) Inhibition of wild type SARS-CoV-2 infection in ACE2-expressing A549 cells by 10 µM amodiaquine. (**c**) Reduction of viral load in the lungs of hamsters treated once a day with amodiaquine (50 mg/kg) beginning 1 day prior to intranasal administration of SARS-CoV-2 virus (10^3^ PFU) as measured by qPCR for subgenomic RNA encoding SARS-CoV-2 N protein. **, p< 0.01. (**d**) Hematoxylin- and SARS-CoV-2 N-stained histological sections of lungs from animals that were mock treated, infected with SARS-CoV-2 and treated with vehicle alone, or infected with SARS-CoV-2 and treated with amodiaquine (50 mg/kg subcutaneously). (**e**) Reduction of viral load in the lungs of hamsters treated once a day for 4 days with amodiaquine (50 mg/kg) beginning 1 day prior to co-caging with SARS-CoV-2 infected animals as measured by qPCR for RNA encoding SARS-CoV-2 N protein. *, p< 0.05. (f) Graph depicting plaque forming units (PFU) per mL of lung homogenate from hamsters pretreated with vehicle or amodiaquine one day prior to being exposed to infected animals. Each cohort was comprised of 4 animals; p-value = 0.037.

We then used the Huh-7 cells in a 96-well plate assay format to test the effects of multiple drugs on SARS-CoV-2pp entry that have been approved by the FDA for other medical indications, including chloroquine, hydroxychloroquine, amodiaquine, toremifene, clomiphene, arbidol, verapamil, and amiodarone. These drugs were chosen based on the hypothesis that they might have broad-spectrum antiviral activity because they have been shown to inhibit infection by other SARS, influenza, and Ebola viruses^31–33^. All of these drugs demonstrated dose-dependent inhibition of SARS-CoV-2pp entry in Huh-7 cells without producing any detectable cell toxicity (**Extended Data Fig. 6)** when added at 1 and 5 µM simultaneously with the virus and culturing for 72 hours (**Fig. 4a**). These results were promising; however, Huh-7 cells only express low levels of ACE2^34^ and they do not express TMPRSS2^29, 35^. In addition, this cell line was derived from a human liver tumor, whereas SARS-CoV-2 preferentially targets lung in humans.

Thus, to test the clinical translation potential of the drugs that were active in the Huh-7 cell assay, we evaluated their ability to prevent SARS-CoV-2pp infection in the more highly differentiated and physiologically relevant human Lung Airway Chips. SARS-CoV-2pp were introduced into the air channel of the Airway Chips to mimic human infection by airborne SARS-CoV-2. High levels of the viral pol gene encoded by the SARS-CoV-2pp were detected in the lung airway epithelial cells in chips infected by SARS-CoV-2pp within 48 hours, but not in control chips that were inoculated with pseudoparticles without SARS-CoV-2 spike protein (**Extended Data Fig. 7a**). Infection with SARS-CoV-2pp was also blocked by a neutralizing antibody that targets the receptor binding domain (RBD) of SARS-CoV-2 (**Extended Data Fig. 7b**), confirming that entry of the pseudotyped SARS-CoV-2 virus into the epithelial cells of the human Lung Airway Chip is mediated specifically by the SARS-CoV-2 S protein. The ability of SARS-CoV-2pp to efficiently infect human airway epithelial cells on-chip is consistent with our finding that these highly differentiated lung cells express high levels of its ACE2 receptor as well as TMPRSS2 (**Fig. 1c-e**), which mediate cellular entry of native SARS-CoV-2 virus^18, 19^. In addition, immunofluorescence microscopic analysis confirmed that the SARS-CoV-2pp preferentially infected ciliated cells in the human Lung Airway Chip (**Extended Data Fig. 7c**), as native SARS-CoV-2 virus does in vivo^19^.

Next, we pretreated the human Airway Chips by perfusing their vascular channel for 24 hours with amodiaquine, toremifene, clomiphene, chloroquine, hydroxychloroquine, arbidol, verapamil, or amiodarone at clinically relevant levels similar to their maximum concentration (C_max_) in blood reported in humans (**Table 1**) to mimic systemic distribution after oral administration. SARS-CoV-2pp were then introduced into the airway channel and incubated statically while continuously flowing the drug through the vascular channel for additional 48 hours. qPCR quantitation of viral mRNA revealed that only three of these drugs — amodiaquine, toremiphene, and clomiphene — significantly reduced viral entry (by 59.1%, 51.1% and 28.1%, respectively) (**Fig. 4b**) without producing detectable cytotoxicity (**Extended Data Fig. 6b**) under these more clinically relevant experimental conditions. Importantly, hydroxychloroquine, chloroquine, and arbidol that have no effect on SARS-CoV-2pp entry in our human Airway Chips also failed to demonstrate clinical benefits in human clinical trials^1,2,36^. When administered to patients, the most potent drug amodiaquine is rapidly transformed (half life ∼ 5 hr) into its active metabolite, desethylamodiaquine, which has a much longer half-life (∼ 9-18 days)^37^. When desethylamodiaquine was administered at a clinically relevant dose (1 µM; **Table 1**) in the human Airway Chips, it also reduced entry of the pseudotyped SARS-CoV-2 viral particles by ∼60% (**Extended Data Fig. 8**) suggesting that both amodiaquine and its metabolite are active inhibitors of SARS-CoV-2 S protein-dependent viral entry.

**Table 1.** Clinically relevant drug concentrations used in human Airway Chips.

| Drug | C<sub>max</sub> (ng/ml) | C<sub>max</sub> (μM) |
| --- | --- | --- |
| Amodiaquine | 575 | 1.24 |
| Toremifene | 1211 | 2.98 |
| Clomiphene | 500 | 0.83 |
| Chloroquine | 960.5 | 1.91 |
| Hydroxychloroquine | 422 | 1.25 |
| Arbidol | 2160 | 3.89 |
| Verapamil | 287 ± 105 | 0.81 |
| Amiodarone | 13660 ± 3410 | 20.04 |
| Desethylamodiaquine | 329-828 | 1.00 |

## Amodiaquine and desethylamodiaquine inhibit SARS-CoV-2 infection *in vitro* and *in vivo*

Finally, we tested the ability of the most potent drug identified in the Airway Chip, amodiaquine, and its metabolite desethylamodiaquine to inhibit infection by GFP-labeled SARS-CoV-2 virus at a multiplicity of infection (MOI = 0.1) in Vero E6 cells. We found that both compounds inhibited infection by SARS-CoV-2 in a dose-dependent manner (**Fig. 5a**) with half maximal inhibitory concentrations (IC_50_) of 10.3 ± 1.6 and 8.5 ± 3.0 µM for amodiaquine and desethylamodiaquine, respectively. Amodiaquine and desethylamodiaquine also inhibited infection by wild type SARS-CoV-2 virus when administered under less stringent conditions (MOI = 0.01), with both compounds exhibiting IC_50_ < 5 µM. In addition, amodiaquine reduced viral load by ∼ 3 logs in ACE2-expressing human lung A549 cells infected with native SARS-CoV-2 when administered at 10 µM (**Fig. 5b**).

Given this potent inhibitory activity against native SARS-CoV-2, we then evaluated amodiaquine in a hamster COVID-19 model in which the animals are infected intranasally with SARS-CoV-2 virus (10^3^ PFU). The animals were treated once a day for 4 days with amodiaquine (50 mg/kg via subcutaneous injection) beginning one day prior to SARS-CoV-2 infection. The dosing regimen was selected based on a PK study for amodiaquine that was carried out in healthy hamsters in parallel. A single dose of amodiaquine (50 mg/kg) injected subcutaneously revealed that the C_max_ for amodiaquine and its active metabolite desethylamodiaquine were ∼ 3.2 and 0.7 µM, respectively; while the T_1/2_ for amodiaquine was 18.1 hours, that of its active metabolite was significantly greater than the 1 day timecourse analyzed (**Extended Data Fig. 9a,b**), which is consistent with human clinical data^37^. Analysis of drug concentrations in lung, kidney, intestine, and heart revealed that both drugs became concentrated in these organs relative to plasma (**Extended Data Fig. 9c**). Analysis of drug concentrations 24 hours after dosing revealed significant exposures of amodiaquine and desethylamodiaquine in lung, kidney, and intestine (**Extended Data Fig. 9c**), with levels in tissues relative to plasma enhanced 21- to 138-fold for amodiaquine and 8- to 45-fold for desethylamodiaquine. These PK results, including the extended half lives and tissue concentration for both compounds are consistent with results of past PK studies in humans^37^.

Importantly, amodiaquine treatment of infected hamsters resulted in ∼70% reduction in SARS-CoV-2 viral load measured by RT-qPCR of the viral N transcript when measured 3 days after the viral challenge (**Fig. 5c**). Immunohistochemical analysis of lungs from these animals confirmed that amodiaquine treatment resulted in a significant reduction in expression of SARS-CoV-2 N protein in these tissues (**Fig. 5d**). We then carried out studies using a SARS-CoV-2 transmission model in which vehicle or amodiaquine-treated healthy animals are placed in the same cage with animals that had been infected with SARS-CoV-2 virus one day earlier. In vehicle controls, this experimental setup results in a 100% transmission within two days of exposure. In contrast, the same amodiaquine treatment regimen as described above resulted in a 90% inhibition of SARS-CoV-2 infection as measured by quantifying N transcript levels (**Fig. 5e**). These results were further corroborated in an independent experiment where amodiaquine-treated animals showed a greater than one log decrease in viral titers measured by plaque assays when compared to vehicle (**Fig. 5f**). To our knowledge, this is the first example of successful chemoprophylaxis against SARS-CoV-2 *in vivo*. Taken together, these results confirm that the antiviral activities identified in the human Lung Airway Chips translate to the *in vivo* setting, and suggest that amodiaquine may provide significant protection when taken prophylactically.

## Discussion

Taken together, these data show that human Organ Chips, such as the Lung Airway Chip, can be used to rapidly identify existing approved drugs that may be repurposed for pandemic virus applications in crisis situations that require accelerated development of potential therapeutic and prophylactic interventions. Our work on repurposing of therapeutics for COVID-19 was initiated on January 13, 2020 (1 day after the sequence of viral genome was published in GenBank^28^), and our first results with drugs in Airway Chips were obtained three weeks later. The ability to apply drugs using dynamic fluid flow on-chip enables the human lung cells to be treated with more clinically relevant dynamic drug exposures on-chip. While drugs were administered at levels similar to their C_max_ here to compare relative potencies, one caveat is that we did not quantify drug absorption or protein binding in this study. Importantly, by carrying out mass spectrometry measurements of drug levels in these devices, full PK profiles can be recapitulated in these Organ Chip models^8^, which should further aid clinical translation in the future. While animal models remain the benchmark for validation of therapeutics to move to humans, it is important to note that human Organ Chips are now being explored as viable alternatives to animal models^38^ and regulatory agencies are encouraging pharmaceutical and biotechnology companies to make use of data from Organ Chips and other microphysiological systems in their regulatory submissions^39^.

These studies led to the identification of multiple approved drugs that could serve as prophylactics and therapeutics against viral pandemics. The anticoagulant drug, nafamostat, significantly extended the current treatment time window of oseltamivir from 2 to 4 days after infection by influenza virus, which could have great clinical relevance given that most patients do not begin treatment until days after they are infected. Similarly, while the human Organ Chip model successfully predicted the inability of chloroquine, hydroxychloroquine, and arbidol to work in animals^4^ and human patients^1, 2, 40^, in contrast with what was reported in cell lines^41, 42^, we successfully identified amodiaquine as a putative therapeutic for SARS-CoV-2 that works both *in vitro* and *in vivo*. Amodiaquine is an anti-malarial drug related to chloroquine and hydroxychloroquine^43^. This drug was the most potent inhibitor of SARS-CoV-2pp entry into human airway cells, producing ∼60% inhibition when administered under flow at 1.24 µM, which should be clinically achievable in the plasma of patients with malaria who receive 300 mg administration^44^, as well as in tissues such as lung where the drug and its metabolite concentrate^37^. Importantly, further investigation of amodiaquine revealed that both this drug and its active metabolite (desethylamodiaquine) do indeed inhibit native SARS-CoV-2 infection *in vitro* and *in vivo*. Thus, these findings suggest that the microfluidic human Organ Chip model, combined with existing preclinical cell-based and animal assays, offers a potentially more clinically relevant test bed for accelerated discovery of anti-COVID-19 drugs.

When considering repurposing of approved drugs for COVID-19, it is important to recognize that every drug has its own distinct therapeutic and toxicity profile that must be taken into consideration. Amodiaquine has been widely used for prophylaxis and treatment of malaria for over 60 years. It is currently used in low resource nations where the World Health Organization (WHO) recommends it be used in combination with artesunate for chemoprophylaxis of malaria and as a second line acute treatment for uncomplicated *P. falciparum*-resistant malaria. Interestingly, the amodiaquine-artesunate drug combination also has been reported to lower the risk of death from Ebola virus disease^45^. But amodiaquine was withdrawn from use in the United States due to rare occurrence of agranulocytosis and liver damage with high doses or prolonged treatment^46^; however, it continues to be well tolerated among African populations where it is commonly administered for short duration (3 day course). The short course is possible because the half life of amodiaquine’s active metabolite, desethylamodiaquine, is very long (on the order of 9 to 18 days) and it concentrates in organs, including lung^37^.

Given the alarming rate at which the SARS-CoV-2 pandemic is spreading, clinicians must seriously consider the relative risks and benefits of using any existing approved drug as a new COVID-19 therapy with specific patient populations (e.g., male versus female, young versus old, Caucasian versus African, etc.) before initiating any trial in their local communities. Our findings raise the possibility that amodiaquine could be explored as a chemoprophylaxis therapy to prevent spread of COVID-19 and help people return to their workplace by treating healthy patients for 3 days, which could then offer protection for an additional 2 weeks. If amodiaquine were to be advanced to clinical trials for prevention of COVID-19, it would be critical to select patient populations carefully and appropriate clinical assessments (e.g., blood and liver function tests) should be carried out before and during administration of drug. This prophylactic therapy may be particularly valuable in Africa and other low resource nations where this inexpensive drug is more readily available and where more expensive alternative therapies are not feasible.

The current COVID-19 pandemic and potential future ones caused by influenza viruses or other coronaviruses, represent imminent dangers and major ongoing public health concerns. When it comes to repurposing existing antiviral agents, every experimental assay has its limitations. However, our results suggest that combining multiplexed cell-based assays with lower throughput/higher content human Organ Chips that recapitulate human-relevant responses as well as animal models, and focusing on compounds that are active in all models, could provide a fast track to identify potential treatments for the current COVID-19 pandemic that have a higher likelihood of working in human patients. This discovery pipeline may be equally valuable to combat unforeseen biothreats, such as new pandemic influenza or coronavirus strains, in the future.

## METHODS

### Human Airway Chip Culture

Microfluidic two-channel Organ Chip devices containing membranes with 7 μm pores were obtained from Emulate Inc. (Boston, MA). Each microdevice contains two adjacent parallel microchannels (apical, 1 mm wide × 1 mm high; basal, 1 mm wide × 0.2 mm high; length of overlapping channels, 16.7 mm) separated by the porous membrane. Similar results were also obtained in some studies not involving immune cell recruitment using 2-channel devices fabricated from poly-dimethyl siloxane with a PET membrane containing 0.4 um pores, as used in past Airway Chip studies^16^. Before cell plating, both channels of these devices were washed with 70% ethanol, filled with 0.5 mg/mL ER1 solution in ER2 buffer (Emulate Inc.) and placed under UV lamp (Nailstar, NS-01-US) for 20 min to activate the surface for protein coating. The channels were then washed sequentially with ER2 buffer and PBS. The porous membranes were coated on both sides with collagen type IV from human placenta (0.5 mg/mL in water; Sigma-Aldrich) at room temperature overnight. The solution was then aspirated from the chip, which was then used for seeding cells.

Primary human lung airway epithelial basal stem cells (Lonza, USA; Catalog #: CC-2540S) obtained from healthy donors 448571, 446317, 623950, 485960, and 672447) were expanded in 75 cm^2^ tissue culture flasks using airway epithelial cell growth medium (Promocell, Germany) until 60-70% confluent. Primary human pulmonary microvascular endothelial cells (Cell Biologics, USA) were expanded in 75 cm^2^ tissue culture flasks using human endothelial cell growth medium (Cell Biologics, USA) until 70-80% confluent.

To create the human Airway Chips, endothelial cells (2 × 10^7^ cells/mL) were first seeded in the bottom channel by inverting the chip for 4 h in human endothelial cell growth medium, followed by inverting the chip again and seeding of the top channel with the lung airway epithelial basal stem cells (2.5 × 10^6^ cells/mL) for 4 h in airway epithelial cell growth medium. The respective medium for each channel was refreshed and the chips were incubated under static conditions at 37°C under 5% CO_2_ overnight. The adherent cells were then continuously perfused with the respective cell culture medium using an IPC-N series peristaltic pump (Ismatec) or Zoe (Emulate) at a volumetric flow rate of 60 µL/h. After 5-7 days, the apical medium was removed while allowing air to fill the channel to establish an ALI, and the airway epithelial cells were cultured for 3-4 additional weeks while being fed only by constant flow of PneumaCult-ALI medium (StemCell) supplemented with 0.1% VEGF, 0.01% EGF, and 1mM CaCl_2_ from an Endothelial Cell Medium Kit (Cell Biological, M1168) through the bottom vascular channel. The chips were cultured in an incubator containing 5% CO_2_ and 16-18% O_2_ at 85-95% humidity, and the apical surface of the epithelium was rinsed once weekly with PBS to remove cellular debris and mucus. Highly differentiated human airway structures and functions can be maintained in the human lung Airway Chip for more than 2 months.

### Immunofluorescence microscopy

Cells were washed with PBS through the apical and basal channels, fixed with 4% paraformaldehyde (Alfa Aesar) for 20-25 min, and then washed with PBS before being stored at 4°C. Fixed tissues were permeabilized on-chip with 0.1% Triton X-100 (Sigma-Aldrich) in PBS for 5 min, exposed to PBS with 10% goat serum (Life Technologies) and 0.1% Triton X-100 for 30 min at room temperature, and then incubated with primary antibodies (**Supplementary Table 2**) diluted in incubation buffer (PBS with 1% goat serum and 0.1% Triton X-100) overnight at 4°C, followed by incubation with corresponding secondary antibodies (**Supplementary Table 2**) for 1 h at room temperature; nuclei were counterstained with DAPI (Invitrogen) after secondary antibody staining. Fluorescence imaging was carried out using a confocal laser-scanning microscope (SP5 X MP DMI-6000, Germany) and image processing was done using Imaris software (Bitplane, Switzerland).

### Barrier function assessment

To measure tissue barrier permeability, 50 µl cell medium containing Cascade blue (607 Da) (50 µg/mL; Invitrogen) was added to bottom channel and 50 µl cell medium was added to top channel. The fluorescence intensity of medium of top and bottom channels was measured 2 h later in three different human Airway chips. The apparent permeability was calculated using the formula: P_app_ = *J*/(*A* × ΔC), where P_app_ is the apparent permeability, *J* is the molecular flux, *A* is the total area of diffusion, and ΔC is the average gradient.

### Mucus quantification

Mucus present in the airway channel was isolated by infusing 50 µl PBS into the upper channel of the Airway Chip, incubating for 1 h at 37°C, and then collecting the fluid and storing it at −80°C before analysis, as previously described^6^. Quantification of mucus production was carried out by quantifying Alcian Blue Staining (Thermo Fisher Scientific) and comparing to serially diluted standards of mucin (Sigma-Aldrich) in PBS.

### Quantitative reverse transcription-polymerase chain reaction (RT-qPCR)

Total RNA was extracted from differentiated human Airway chips, pre-differentiated lung airway epithelial cells, or MDCK cells using TRIzol (Invitrogen). cDNA was then synthesized using AMV reverse transcriptase kit (Promega) with Oligo-dT primer. To detect cellular gene-expression level, quantitative real-time PCR was carried out according to the GoTaq qPCR Master Mix (Promega) with 20 µl of a reaction mixture containing gene-specific primers (**Supplementary Table 3**). The expression levels of target genes were normalized to GAPDH.

### Influenza viruses

Influenza virus strains used in this study include A/PR/8/34 (H1N1), GFP-labeled A/PR/8/34 (H1N1), A/WSN/33 (H1N1), A/Netherlands/602/2009 (H1N1), A/Hong Kong/8/68/ (H3N2), A/Panama/2007/99 (H3N2), and A/Hong Kong/156/1997 (H5N1). A/PR/8/34 (H1N1), GFP-labeled A/PR/8/34 (H1N1), A/WSN/33 (H1N1) were generated using reverse genetics techniques. Other viruses were obtained from the Centers for Disease Control and Prevention (CDC) or kindly shared by Drs. P. Palese, R.A.M. Fouchier, and A. Carcia-Sastre.

### Influenza virus infection of human Airway Chips

Human Airway Chips were infected with influenza viruses by flowing 30 uL of PBS containing the indicated multiplicity of infection (MOI) of viral particles into the apical channel, incubating for 2 h at 37°C under static conditions, and then removing the medium to reestablish an ALI. To measure virus propagation, the apical channel was incubated with 50 µl of PBS for 1 h at 37°C at various times, and then the apical fluid and vascular effluent were collected from the apical and basal channels, respectively, to quantify viral load using the plaque formation assay; released cytokines and chemokines were analyzed in these same samples. The tissues cultured on-chip were also fixed and subjected to immunofluorescence microscopic analysis.

To test the efficacy of oseltamivir acid, Airway Chips infected with influenza virus (MOI = 0.1) were treated with 1 µM oseltamivir acid (Sigma-Aldrich) under flow (60 µl/h) through the vascular channel. To explore the effects of serine protease inhibitors on influenza infection, Nafamostat (Abcam) or Trasylol (G-Biosciences) was delivered into the airway channel of influenza-infected chip (MOI = 0.1). Two days later, the virus samples were collected for detection of viral load and the vascular effluents were collected for analysis of cytokines and chemokines. In the treatment time window detection experiment, oseltamivir acid (1 µM), nafamostat (10 µM), or both were added to the influenza H1N1-infected Airway Chips (MOI = 0.1) at indicated times. Oseltamivir was perfused through the vascular channel, while nafamostat was introduced in 20 uL of PBS and incubated in the airway channel for 48 hours. Fluids samples were then collected from both channels for detection of viral load.

### Analysis of neutrophil infiltration

Neutrophils isolated from fresh human blood using a Ficoll-Paque PLUS (GE Healthcare) gradient were resuspended in medium at a concentration of 5 × 10^6^ cells/mL, which is within the normal range (2.5-7.5 × 10^6^ cells/ml) of neutrophils found in human blood. The isolated neutrophils were labeled with Cell Tracker Red CMTPX (Invitrogen) and injected into the vascular channel of inverted Airway Chips infected with influenza virus (MOI = 0.1) at a flow rate of 50-100 µL/h using a syringe pump; 2 h later unbound neutrophils were washed away by flowing cell-free medium for 24 h. Virus samples were collected by incubating the airway channel with 50 µl of PBS for 1 h at 37°C, collecting the fluid, and detecting virus load using the plaque assay. The cell layers were fixed on-chip and subjected to immunofluorescence microscopic analysis for influenza virus NP (Invitrogen) and neutrophils (CD45, Biolegend). Micrographs of four or five random areas were taken from chips for subsequent quantification of infiltrated neutrophils. To study the interaction between influenza virus and neutrophils, Airway Chips were infected with GFP-labeled PR8 virus (MOI = 0.1) for 24 h. Cell Tracker Red CMTPX-labeled neutrophils (5 × 10^6^ cells/mL) were perfused in medium through the vascular channel of infected Airway Chips. Immunofluorescence microscopic analysis were carried out at indicated times.

### Plaque formation assay

Virus titers were determined using plaque formation assays. Confluent MDCK cell monolayers in 12-well plate were washed with PBS, inoculated with 1 mL of 10-fold serial dilutions of influenza virus samples, and incubated for 1 h at 37°C. After unabsorbed virus was removed, the cell monolayers were overlaid with 1 mL of DMEM (Gibco) supplemented with 1.5% low melting point agarose (Sigma-Aldrich) and 2 µg/mL TPCK-treated trypsin (Sigma-Aldrich). After incubation for 2-4 days at 37°C under 5% CO_2_, the cells were fixed with 4% paraformaldehyde, and stained with crystal violet (Sigma-Aldrich) to visualize the plaques; virus titers were determined as plaque-forming units per milliliter (PFU/mL). Plaque titers from in vivo lung samples were determined post mortem by complete lung dissection and dissociation in PBS. Debris was pelleted a 5000 rpm and the remaining supernatant was used to determine PFU/mL.

### Analysis of cytokines and chemokines

Vascular effluents from Airway Chips were collected and analyzed for a panel of cytokines and chemokines, including IL-6, IP-10, MCP-1, RANTES, interferon-β, using custom ProcartaPlex assay kits (Invitrogen). Analyte concentrations were determined using a Luminex100/200 Flexmap3D instrument coupled with Luminex XPONENT software (Luminex, USA).

### Analysis of cleavage of virus hemagglutinin (HA) by serine proteases

For analysis of HA cleavage by serine proteases in the presence or absence of nafamostat, MDCK cells (5 × 10^5^ cells per well in 6-well plates) were transfected with 2·5 µg serine protease expression plasmid or empty vector using TransIT-X2 Dynamic Delivery System (Mirus). One day later, the cells were infected with influenza A/WSN/33 (H1N1) virus (MOI = 0·01) in DMEM supplemented with 1% FBS, and then cultured in the presence or absence of 10 µM nafamostat. Two days post-infection, the supernatant was harvested and subjected to Western blot analysis using anti-HA1 antibody.

### Drugs for the SARS-CoV2pp studies

Chloroquine (cat. #ab142116), Hydroxychloroquine (cat. #ab120827), arbidol (cat. #ab145693), toremifene (cat. #ab142467), clomiphene (cat. #ab141183), verapamil (cat. #ab146680), and amiodarone (cat. #ab141444) were purchased from Abcam; amodiaquine dihydrochloride dihydrate (cat. #A2799) was purchased from Sigma-Aldrich; N-desethylamodiaquine (cat. #20822) was purchased from Caymanchem. Chloroquine was dissolved in water to a stock concentration of 10 mM; all other tested drugs were dissolved in dimethyl sulfoxide (DMSO) to a stock concentration of 10 mM. The purity of all evaluated drugs was > 95%.

### Plasmids

Plasmid expressing the spike protein of SARS-CoV-2 (pCMV3-SARS-CoV2-Spike) was purchased from Sino Biological Inc. (Beijing, China). pCMV-VSVG, pNL4-3.Luc.R-E-, and pAdvantage were obtained from Addgene, NIH AIDS Reagent Program, and Promega, respectively. All plasmids used for transfection were amplified using the Maxiprep Kit (Promega) according to the manufacturer’s instructions.

### Pseudotyped virus production

HEK293T cells (5 × 10^5^ cell per well) were seeded into 6-well plates. 24 h later, HEK293T cells were transfected with 1.0 µg of pNL4-3.Luc.R-E-, 0.07 µg of pCMV3-SARS-CoV-2-Spike, and 0.3 µg of pAdvantage with the TransIT-X2 transfection reagent (Mirus) according to the manufacturer’s instructions to produce SARS-CoV-2 spike pseudotyped HIV virions (SARS-CoV-2pp). Similarly, HEK293T cells were transfected with 1.0 µg of pNL4-3.Luc.R-E-, 0.7 µg of pCMV-VSVG, and 0.3 µg of pAdvantage to produce VSVG pseudotyped HIV virions (VSVpp). The supernatants containing the pseudotyped viruses were collected at 48 h post-transfection and clarified by the removal of floating cells and cell debris with centrifugation at 10^3^ g for 5 min. The culture supernatants containing pseudotyped viruses particles were either used immediately or flash frozen in aliquots and stored at 80°C until use after being concentrated using a PEG virus precipitation kit (Abcam). Incorporation of the SARS-CoV-2 S protein into the SARS-CoV-2pp was confirmed using Western Blot analysis with anti-SARS-CoV-2 S1 chimeric monoclonal antibody with combined constant domains of the human IgG1 molecule and mouse variable regions (40150-D001 Sinobiological, 1:500); a recombinant receptor binding domain (RBD) fragment from the S1 region was used as a control (BEI resources, NR-52306). Similar results were also obtained using a commercially available pseudotyped SARS-CoV-2 S protein expressing viral particles (Amsbio LLC).

### Infection assay using pseudotyped viruses in Huh-7 cells

Drugs were tested using entry assays for SARS-CoV-2pp and VSVpp, as previously described^27^. Infections were performed in 96-well plates. SARS-CoV-2pp or VSVpp was added to 5 × 10^3^ Huh-7 cells (a human liver cell line) per well in the presence or absence of the test drugs or compounds. The mixtures were then incubated for 72 hours at 37°C. Luciferase activity, which reflects the number of pseudoparticles in the host cells, was measured at 72 h post-infection using the Bright-Glo reagent (Promega) according to the manufacturer’s instructions. Test drugs were serially diluted to a final concentration of 1 or 5 μM. The maximum infectivity (100%) was derived from the untreated wells; background (0%) from uninfected wells. To calculate the infection values, the luciferase background signals were subtracted from the intensities measured in each of the wells exposed to drug, and this value was divided by the average signals measured in untreated control wells and multiplied by 100%.

### SARS-CoV-2pp infection of human lung Airway Chips

To measure infection in human Airway Chips with the pseudotyped virus, drugs were flowed through the vascular channel of the Airway Chips at their reported C_max_ in human blood (**Table 2**) while airway channel was statically treated with the same concentrations of drugs. 24 h later, the SARS-CoV-2pp was delivered into the airway channel in a small volume (30 μL) of medium containing the drug at the same concentrations and incubated statically for additional 48 h while the drug at the same dose was continuously flowed through the vascular channel at 37°C. The lung airway epithelium was then collected by RNeasy Micro Kit (Qiagen) according to the manufacturer’s instructions and subjected to analysis of viral load by qRT-PCR. As we only focused in assessing viral entry in these studies, the chips were only lined by differentiated airway epithelium and did not contain endothelium.

### Native SARS-CoV-2 in vitro infection assay

All work with native SARS-CoV-2 virus was performed in a Biosafety Level 3 laboratory and approved by our Institutional Biosafety Committee. Vero E6 cells (ATCC# CRL 1586) were cultured in DMEM (Quality Biological), supplemented with 10% (v/v) heat inactivated fetal bovine serum (Sigma), 1% (v/v) penicillin/streptomycin (Gemini Bio-products), and 1% (v/v) L-glutamine (2 mM final concentration, Gibco) (Vero Media). Cells were maintained at 37°C (5% CO2). GFP-labeled native SARS-CoV-2 was generously provided by Dr. Ralph S. Baric^47^. Stocks were prepared by infection of Vero E6 cells for two days when CPE was starting to become visible. Media were collected and clarified by centrifugation prior to being aliquoted for storage at −80°C. Titer of stock was determined by plaque assay using Vero E6 cells. GFP-labeled native SARS-CoV-2 infection and drug testing were performed in Vero E6 cells 1. Cells were plated in clear bottom, black 96-well plates one day prior to infection. Drug was diluted from stock to 50 μM and an 8-point 1:2 dilution series prepared in duplicate in Vero Media. Each drug dilution and control was normalized to contain the same concentration of drug vehicle (e.g., DMSO). Cells were pre-treated with drug for 2 h at 37°C (5% CO2) prior to infection with SARS-CoV-2 at MOI = 0.1. Plates were then incubated at 37°C (5% CO2) for 48 h, followed by fixation with 4% PFA, nuclear staining with Hoechst (Invitrogen), and data acquisition on a Celigo 5-channel Imaging Cytometer (Nexcelom Bioscience, Lawrence, CA). The percent of infected cells was determined for each well based on GFP expression by manual gating using the Celigo software. In addition to plates that were infected, parallel plates were left uninfected to monitor cytotoxicity of drug alone. Plates were incubated at 37°C (5% CO2) for 48 h before performing CellTiter-Glo (CTG) assays as per the manufacturer’s instruction (Promega, Madison, WI). Luminescence was read on a BioTek Synergy HTX plate reader (BioTek Instruments Inc., Winooski, VT) using the Gen5 software (v7.07, Biotek Instruments Inc., Winooski, VT). Similar results were obtained with wild type SARS-CoV-2 virus, using a previously published method^45^.

### Hamster PK studies

Amodiaquine dihydrochloride dihydrate (Sigma, #A2799) was formulated at 10 mg/ml in 12% sulfobutylether-β-cyclodextrin in water at pH 5.0 and administered to LVG male hamsters (n=3) at 50 mg/kg by subcutaneous injection (dose volume of 5 ml/kg). Blood samples were drawn at 0.5, 1, 2, 4, 8 and 24 hours and plasma was prepared. At 24 hours, animals were anesthetized and then perfused to clear tissues of blood. Tissues of interest (lung, heart, kidney and intestine) were removed and homogenized at a 1:3 (w/v) ratio in water. The desired serial concentrations of working reference analyte solutions of amodiaquine (Selleckchem) and desethylamodiaquine (Cayman Biochemicals) were achieved by diluting stock solution of analyte with 50% acetonitrile (0.1%Formic acid) in water solution. 20 µL of working solutions were added to 20 μL of the blank LVG hamster plasma to achieve calibration standards of 1 to 1000 ng/mL in a total volume of 40 μL. 40 μL standards, 40 μL QC samples and 40 μL unknown samples (20 µL plasma with 20 µL blank solution) were added to 200 μL of acetonitrile containing internal standard and 0.1% Formic acid mixture for precipitating protein respectively. The samples were then vortexed for 30 s. After centrifugation at 4°C, 3900 rpm for 15 min, the supernatant was diluted 3 times with water. 5 µL of diluted supernatant was injected into the LC/MS/MS system (AB API 5500 LC/MS/MS instrument with a Phenomenex Synergi 2.5µm Polar-RP 100A (50 × 3 mm) column) for quantitative analysis. The mobile phases used were 95% water (0.1% formic acid) and 95% acetonitrile (0.1% formic acid). All PK studies were conducted by Pharmaron in Ningbo, China.

### Hamster Efficacy Studies

SARS-CoV-2 Isolate USA-WA1/ 2020 (NR-52281) was provided by the Center for Disease Control and Prevention. SARS-CoV-2 was propagated in Vero E6 cells in DMEM supplemented with 2% FBS, 4.5 g/L D-glucose, 4 mM L-glutamine, 10 mM Non-Essential Amino Acids, 1 mM Sodium Pyruvate and 10 mM HEPES and filtered through an Amicon Ultracel 15 (100kDa) centrifugal filter. Flow through was discarded and virus resuspended in DMEM supplemented as above. Infectious titers of SARS-CoV-2 stock were determined using a plaque assay in Vero E6 cells in Minimum Essential Media supplemented with 2% FBS, 4 mM L-glutamine, 0.2% BSA, 10 mM HEPES and 0.12% NaHCO3 and 0.7% agar.

3-5 week-old Syrian hamsters were acclimated to the CDC/USDA-approved BSL-3 facility of the Global Health and Emerging Pathogens Institute at the Icahn School of Medicine at Mount Sinai for 2-4 days. In our direct infection model, hamsters were given a subcutaneous injection posteriorly with drug within 2 hours of drug (amodiaquine) reconstitution one day before SARS-CoV-2 infection and every day thereafter until terminal lung harvest on day 3 post infection. Amodiaquine was reconstituted in 12% sulfobutylether-β-cyclodextrin (Selleckchem) in water(w/w) (with HCl/NaOH) at pH 5.0. Hamsters were intranasally infected with 10^3^ PFU of passage 3 SARS-CoV-2 USA-WA1/2020 in 100 µl of PBS and sacrificed on day 3 of infection. Animals were anesthetized by intraperitoneal injection of 200 µl of ketamine and xylazine (4:1) and provided thermal support while unconscious. Whole lungs were harvested and homogenized in 1 mL of PBS, and homogenates were then spun down at 10,000 rcf for 5 minutes; the supernatant was subsequently discarded, and the lung pellet was resuspended in Trizol. The same protocol was used in our animal-to-animal infection model, except amodiaquine was administered to healthy hamsters for one day before they were housed with untreated hamsters that were infected with SARS-CoV-2 one day earlier, and drug continued to be administered daily for 3 more days, after which infection transmission was quantified.

Lung RNA was extracted by phenol chloroform extraction and DNase treatment using DNA-free DNA removal kit (Invitrogen). After cDNA synthesis of RNA samples by reverse transcription using SuperScript II Reverse Transcriptase (invitrogen) with oligo d(T) primers, quantitative RT-PCR was performed using KAPA SYBR FAST qPCR Master Mix Kit (Kapa Biosystems) on a LightCycler 480 Instrument II (Roche) for subgenomic nucleocapsid (N) RNA (sgRNA) and actin using the following primers: Actin forward primer: 5’-CCAAGGCCAACCGTGAAAAG-3’, Actin reverse primer 5’-ATGGCTACGTACATGGCTGG-3’, N sgRNA forward primer: 5’-CTCTTGTAGATCTGTTCTCTAAACGAAC-3’, N sgRNA reverse primer: 5’-GGTCCACCAAACGTAATGCG-3’ Relative sgRNA levels were quantified by normalizing sgRNA to actin expression and normalizing drug-treated infected lung RNA to vehicle-treated infected controls. All RNA Seq data utilized the Illumina TruSeq Stranded mRNA LP as per the manufacturer’s instructions. Illumina libraries were quantified by Qbit and Agilent Bioanalyzer prior to being run on an Illumina NextSeq500 using a high capacity flow cell. All Raw data was processed as described elsewhere^48^. Raw sequencing data files can be found on NCBI GEO (GSE143613).

### Statistical analysis

All results presented are the result of at least two independent experiments, and if not specified, at least three chips per donor were used in each Organ Chip experiment. Tests for statistically significant differences between groups were performed using a two-tailed Student’s t-test and the Bonferroni correction for multiple hypothesis testing. Differences were considered significant when the *P* value was less than 0.05 (*, P<0.05; **, P<0.01; ***, P<0.001; n.s., not significant). All results are expressed as means ± standard deviation (SD); N ≥ 3 in all studies.

## Data and materials availability

Sharing of materials will be subject to standard material transfer agreements. The nucleotide sequences used in the study have been deposited in GeneBank under accession numbers CY034139.1, CY0334138.1, X17336.1, HE802059.1, CY034135.1, CY034134.1, D10598.1, M12597.1, CY176949.1, CY176948.1, CY176947.1, CY176942.1, CY176945.1, CY176944.1, CY176943.1, CY176946.1, DQ487334.1, DQ487333.1, DQ487335.1, DQ487340.1, DQ487339.1, DQ487337.1, DQ487338.1, and DQ487336.1. Additional data are presented in the Supplementary Materials.

## Contributors

L.S., H.B., and D.E.I. conceived this study, and D.E.I. developed the overall collaborative discovery pipeline. L.S. and H.B. performed and analyzed experiments with other authors assisting with experiments and data analysis. M.B. assisted with cytokine detection assay. W.C., C.O., A.J., A.N., and S.K. assisted with RNA extraction and qRT-PCR. D.Z. and G.G. assisted in the characterization of CoV-2pp. R.K.P. assisted in statistical analysis. R.P. and S.E.G. coordinated experiments and managed the project progress. R.M., D.H., K.O., S.H., T.J., R.A.A. and B.R.t. tested the efficacy of amodiaquine against native SARS-CoV-2 in hamster SARS-CoV-2 infection model. K.C. coordinated the hamster PK studies and assisted in the design of dosing and drug formulation in the hamster efficacy studies. J.L., R.H., M.M., S.W., and M.F. tested the activity of amodiaquine and desethylamodiaquine against native SARS-CoV-2 in Vero E6 cells. L.S., H.B. and D.E.I. wrote the manuscript with all authors providing feedback.

## Declaration of interests

D.E.I. is a founder and holds equity in Emulate Inc., and chairs its advisory board. D.E.I., L. S., R. P., H.B., K H. B., and M. R. are inventors on relevant patent applications hold by Harvard University.

## Acknowledgements

We thank the CDC, Dr. A. Garcia-Sastre, Dr. R.A.M. Fouchier, and Dr. X. Sealens for providing the influenza virus strains and the influenza virus rescue systems. We acknowledge research funding from NIH (NCATS 1-UG3-HL-141797-01 and NCATS 1-UH3-HL-141797-01 to D.E.I.), DARPA under Cooperative Agreements (W911NF-12-2-0036 to D.E.I. and W911NF-16-C-0050 to D.E.I., M.F., and B.tO.). Bill and Melinda Gates Foundation (to D.E.I. and M.F.), Marc Haas Foundation (to B.tO.), and the Wyss Institute for Biologically Inspired Engineering at Harvard University (D.E.I.).

**Correspondence and requests for materials** should be addressed to D.E.I., and to B.tO for issues related to hamster studies.

## EXTENDED DATA FIGURE LEGENDS

**Extended Data Fig. 1.**
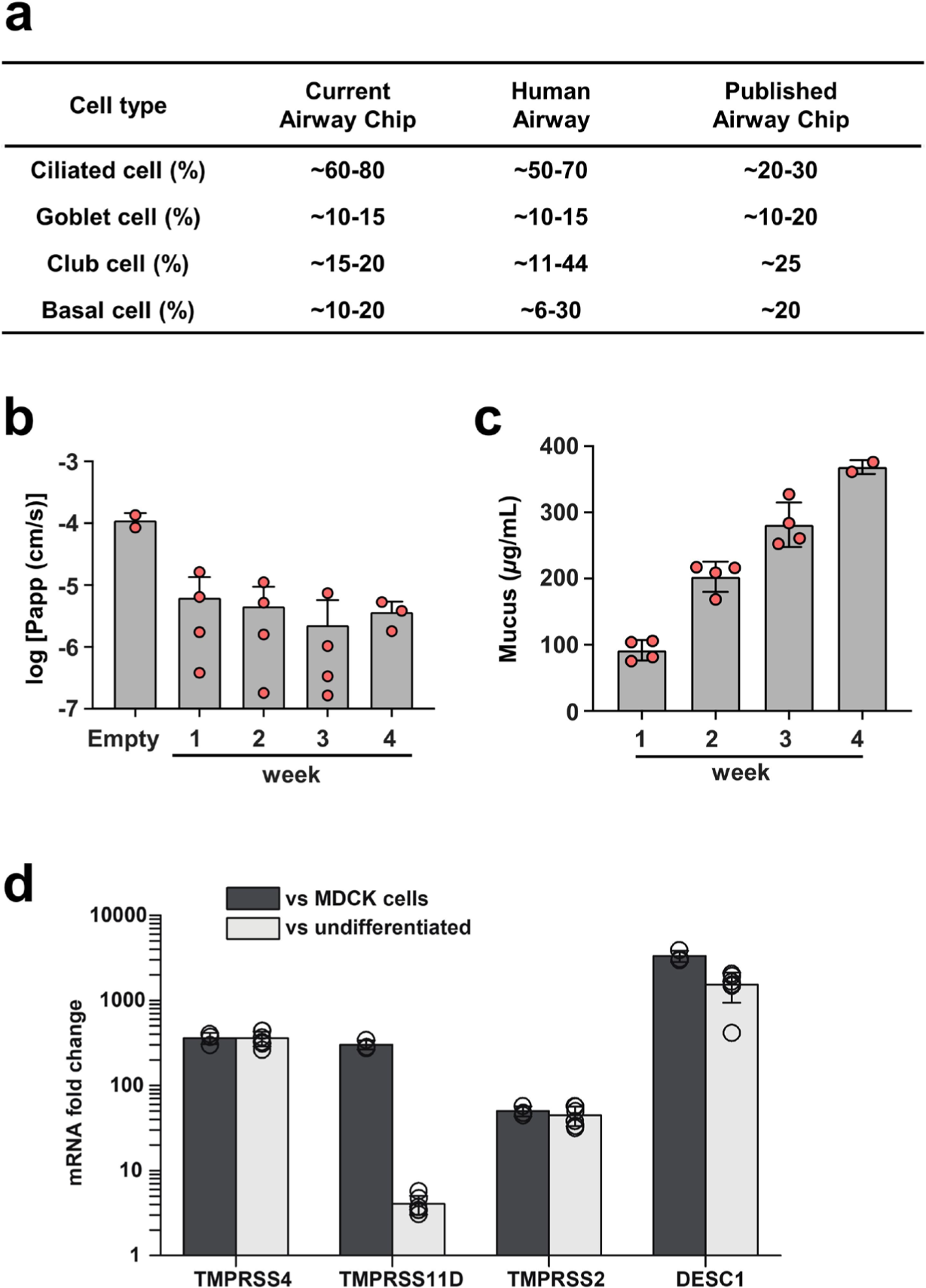
Characterization of human Airway Chip. (**a**) Comparison of the percentage of different lung epithelial cell types in the human Airway Chip presented here compared with those found in living human airway and in our previously published Airway Chip created using a membrane with smaller pores^6, 7^. (**b**) Barrier permeability (log P_app_) of the human Airway Chip assessed using Cascade blue (607 Da) as fluorescent tracer at 1 to 4 weeks of differentiation under an ALI compared with chips without cells (Empty). (**c**) Mucus production at week 1, 2, 3, and 4 post-differentiation quantified using an Alcian Blue assay. (**d**) Fold changes in gene expression levels of 4 different epithelial cell serine proteases (TMPRSS4, TMPRSS11D, TMPRSS2, DESC1) in the well-differentiated Airway Chip versus MDCK cells (one of the most commonly used cell lines in influenza studies) or undifferentiated primary human lung airway epithelial cells.

**Extended Data Fig. 2.**
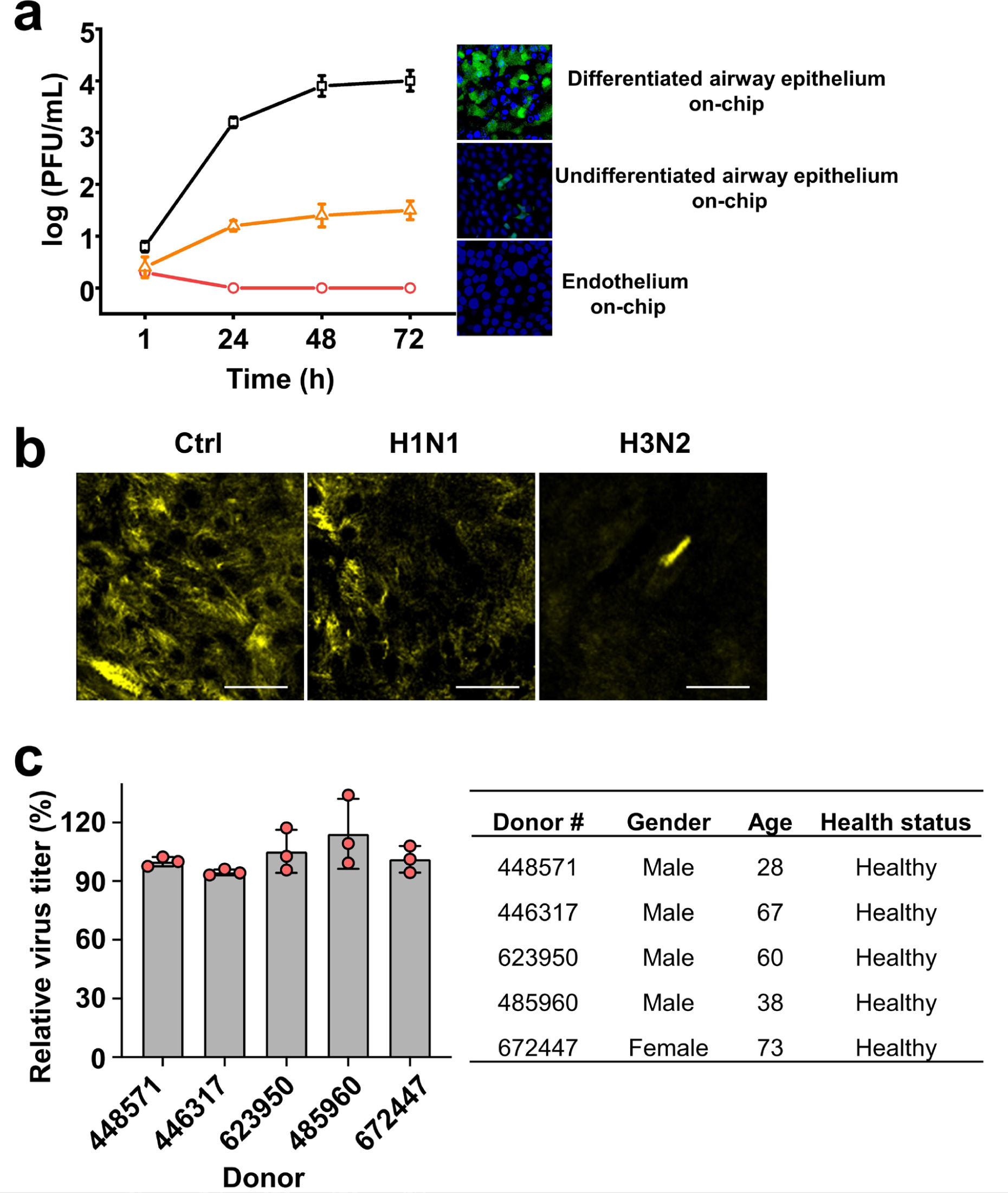
(**a**) Comparison of infectivity and replication of GFP-labeled PR8 (H1N1) in the differentiated epithelium of human lung Airway chip, undifferentiated airway epithelium on-chip, and human vascular endothelium on-chip. Graph showing replication kinetics of influenza PR8 (H1N1) virus (MOI = 0.001) in differentiated epithelium of human Airway Chip, undifferentiated epithelium on-chip, and human vascular endothelium on-chip (left) and corresponding immunofluorescence micrographs showing the infection of GFP-labeled PR8 (H1N1) virus (MOI = 0.1) in these respective chips at 48 h post-infection (green, cells expressing GFP-labeled virus; blue, DAPI-stained nuclei). (**b**) Immunofluorescence micrographs showing apical cilia 24 h post-infection with PR8 (H1N1) or HK/68 (H3N2) (MOI = 0.1) compared to untreated chips (Ctrl). (**c**) Characterization of the replication competence of influenza virus in human lung Airway Chips created with lung airway epithelial basal stem cells obtained from 5 different healthy donors. Influenza PR8 (H1N1) virus was used to infect human Airway chips (MOI = 0.1), and progeny viruses were collected for viral titers detection 48 h later. Information on the donors is shown in the table at the right.

**Extended Data Fig. 3.**
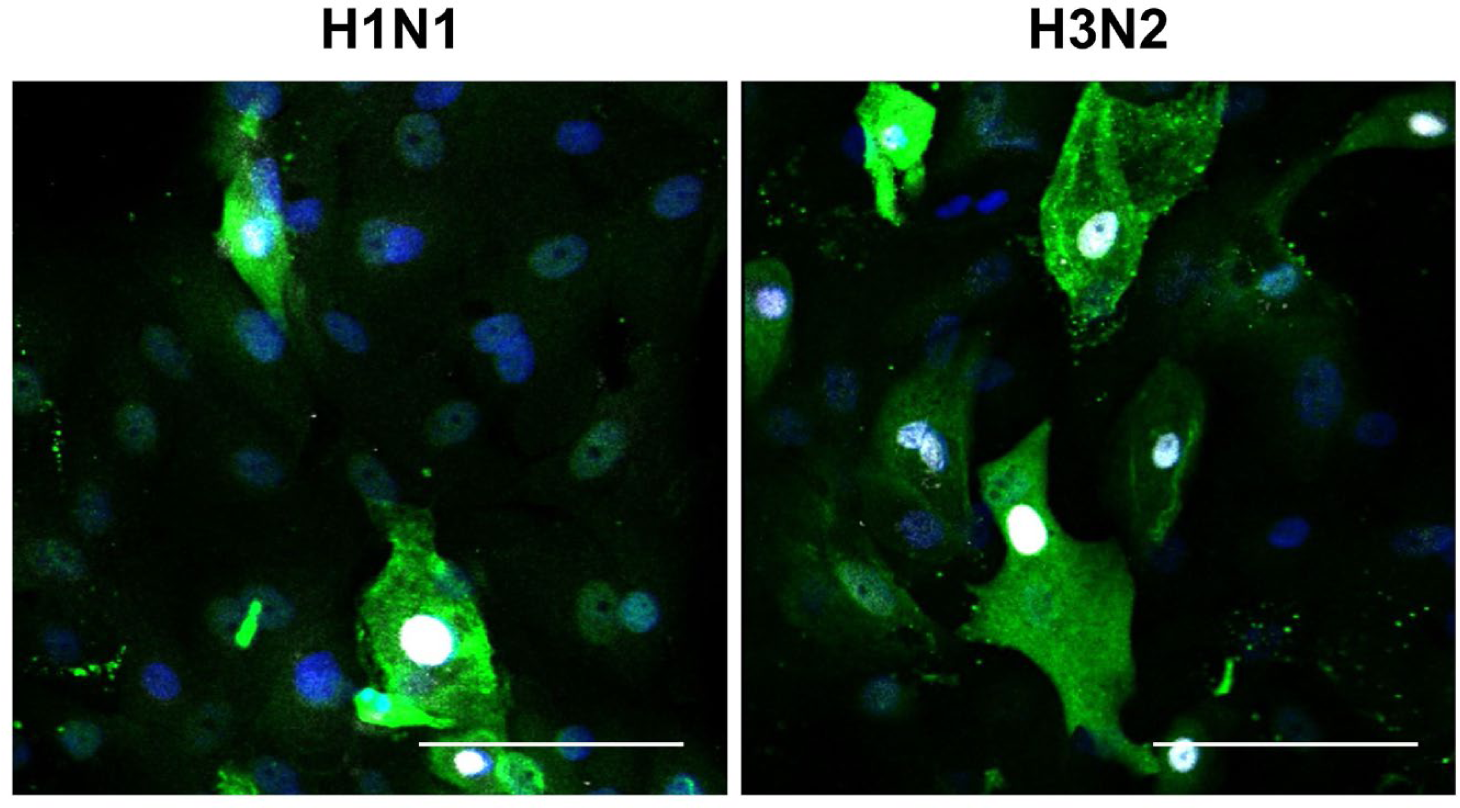
Higher magnification immunofluorescence micrographs showing specific binding of neutrophils (white) to cells infected by WSN (H1N1) or HK/68 (H3N2), which are stained for viral NP (green) (bar, 50 µm).

**Extended Data Fig. 4.**
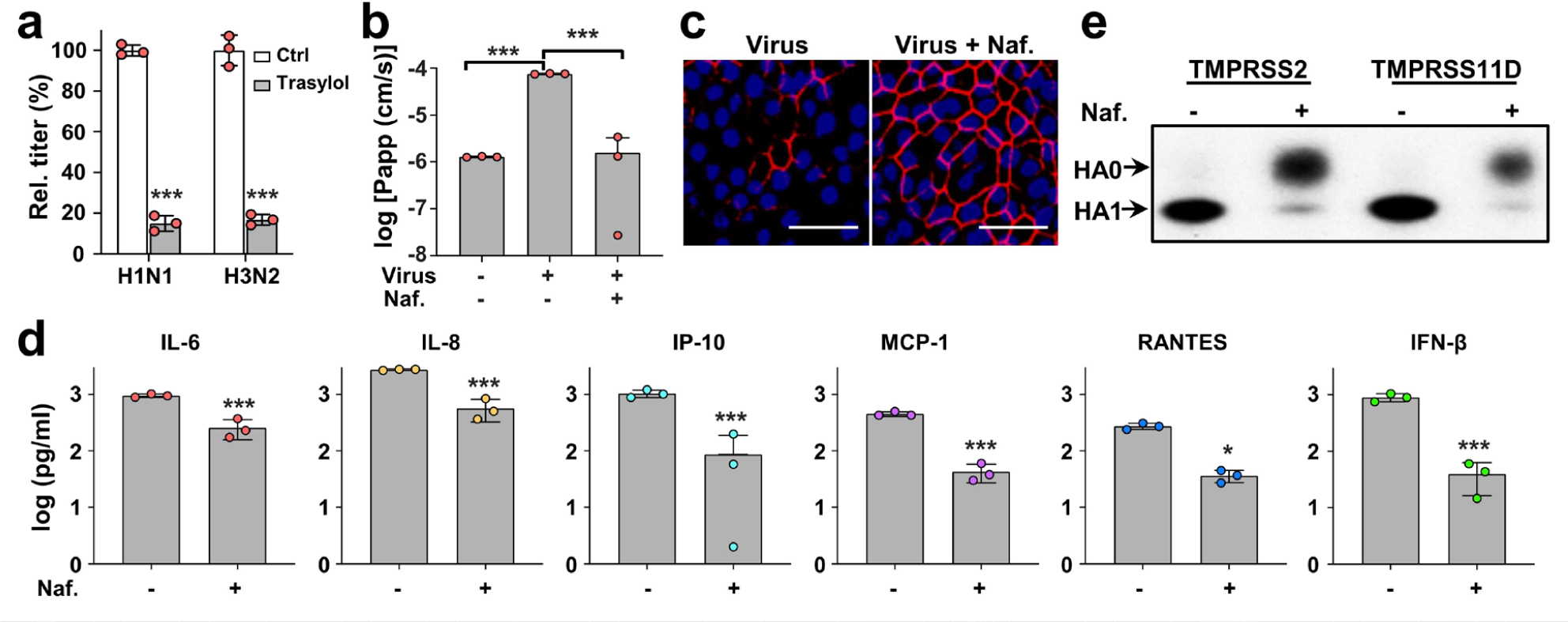
Characterization of FDA-approved protease inhibitor drugs as anti-influenza therapeutics in the human lung Airway Chip. (**a**) Virus titer detection showing the effects of Trasylol (aprotinin) on virus replication of H1N1 and H3N2 in Airway chips 48 h post-infection (Trasylol, gray bars) compared to untreated chips (Ctrl, white bars). (**b**) Barrier permeability (log P_app_) within human Airway Chips measured 48 h post-infection with H1N1 (MOI = 0.1) (+ Virus) in the presence (+) or absence (-) of 10 µM Nafamostat (Naf.) compared to uninfected chips (− Virus). (**c**) Immunofluorescence micrographs showing preservation of ZO1-containing tight junctions seen that are lost in airway epithelium 48 h after infection with H1N1 (MOI = 0.1) (Virus) when treated with 10 µM Nafamostat (Virus + Naf.) (bar, 50 µm). (**d**) Production of various influenza-associated cytokines and chemokines in the human Airway Chip in the presence (+) or absence (-) of 10 µM Nafamostat (Naf.), which suppresses the cytokine response. (**e**) Western blots showing inhibition of TMPRSS2- and TMPRSS11D-mediated cleavage of influenza virus HA0 to HA1 by 10 µM Nafamostat (Naf.).

**Extended Data Fig. 5.**
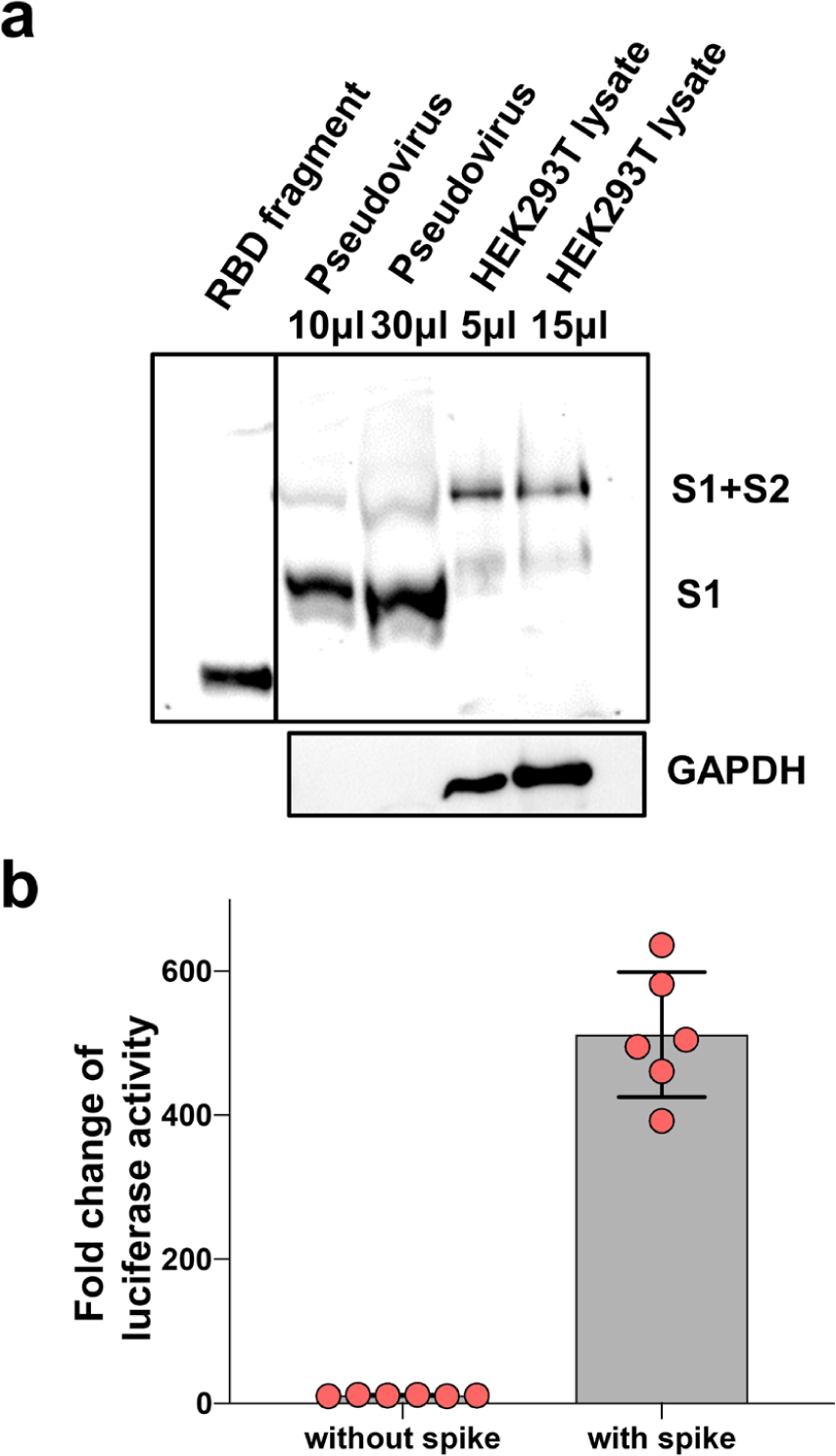
Characterization of the SARS-CoV-2pp and their entry into Huh-7 cells. (**a**) Western blot analysis of SARS-CoV-2 S protein in the lysate of the HEK293T packaging cell line and in pseudotyped virions in the supernatant showing that both uncleaved full-length (S1+S2; ∼180 kDa) and cleaved forms (∼90 kDa) of the spike protein are present in the virions. A recombinant protein containing the receptor binding region domain from S1 (RBD fragment) was used as a positive control, and results were compared to cellular GAPDH. (**b**) Huh-7 cells were infected with SARS-CoV-2pp for 72 h. Luciferase activity was measured to estimate the number of pseudoparticles in the host cells; pseudoparticles without SARS-CoV-2 spike protein were used as control.

**Extended Data Fig. 6.**
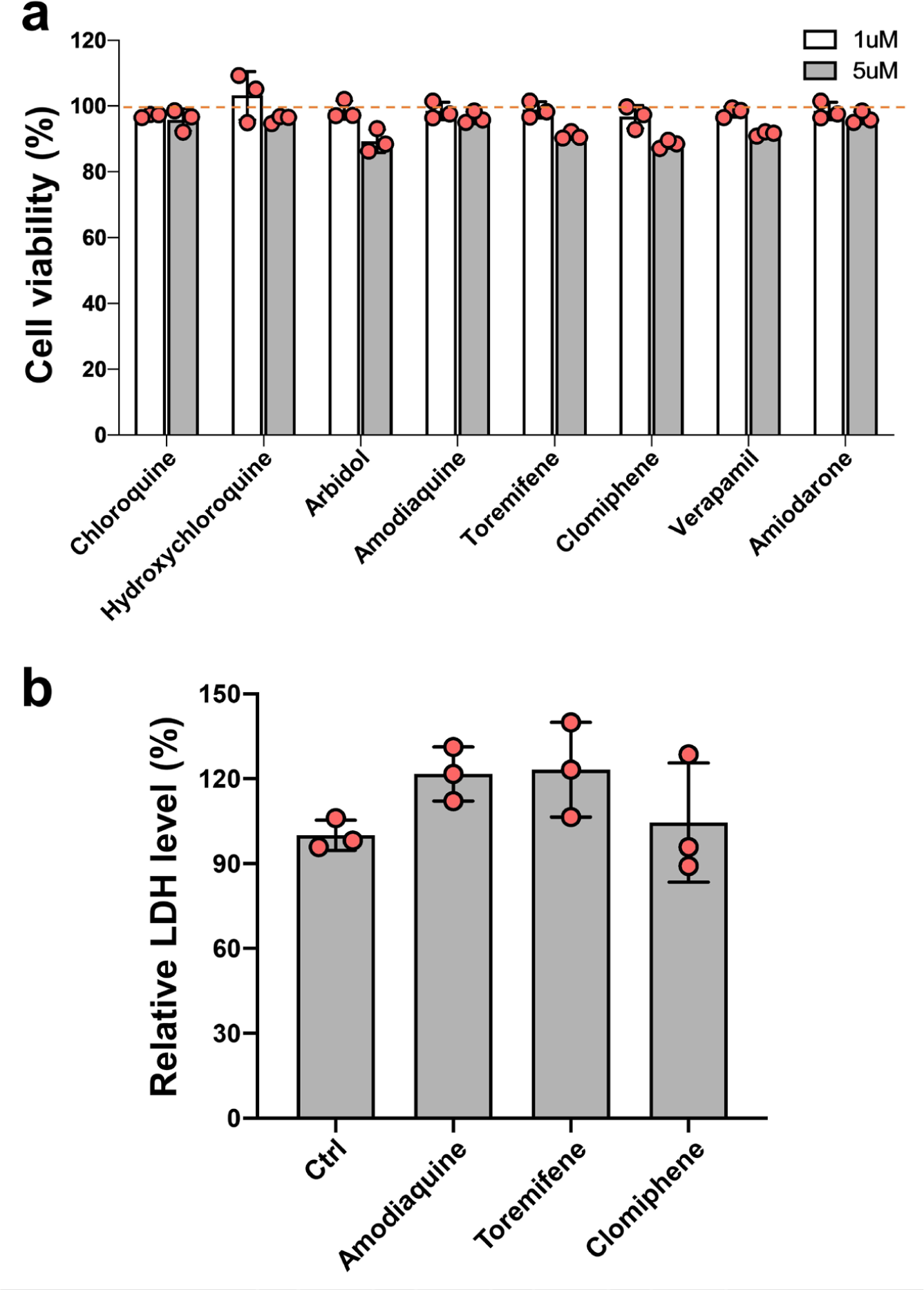
Evaluation of the cytotoxicity of test drugs in Huh-7 cells and human Airway chips. (**a**) Huh-7 cells were treated with the test drugs at 1 or 5 µM for 48 h, and cell viability was measured by Celltiter-Glo assay. The cell viability of untreated cells was set as 100%. (**b**) Human Airway Chips were treated with the test drugs at their respective Cmax for 72 h, cell damage was measure by LDH assay. The LDH level of untreated human Airway Chips was set as 100%. Note that none of the drugs produced any significant cytotoxicity at the doses used in these studies.

**Extended Data Fig. 7.**
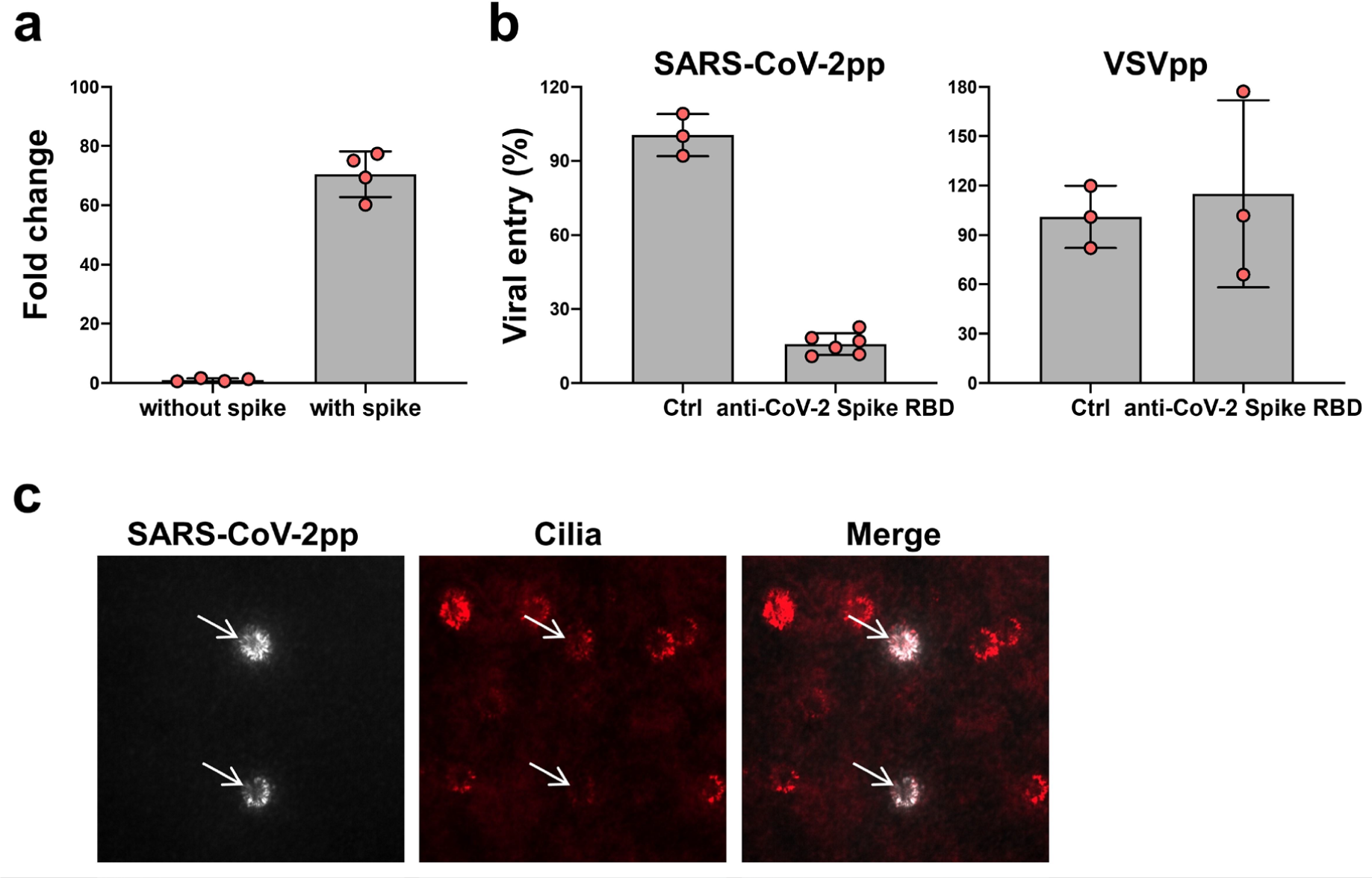
Characterization of SARS-CoV-2pp infection in human Airway Chips. (**a**) SARS-CoV-2 spike protein-mediated infection of CoV-2pp in human Airway Chips. Human Airway Chips were infected with SARS-CoV-2pp for 2 h, washed with PBS, and cultured for 48 h. Cells were collected for detection of viral gene by RT-qPCR. Data are expressed as fold change in viral gene expression relative to control pseudoparticles without SARS-CoV-2 spike protein. (**b**) The infection of SARS-CoV-2pp was blocked by neutralizing antibody targeting the RBD of spike protein. SARS-CoV-2pp or VSVpp were incubated with neutralizing antibody for 1 h at room temperature before infecting human airway epithelium. 48 h later, cells were collected for detection of viral gene by RT-qPCR. PBS was used as control (Ctrl). (**c**) Immunofluorescence micrographs showing specific infection of SARS-CoV-2pp (white) in ciliated cells (green).

**Extended Data Fig. 8.**
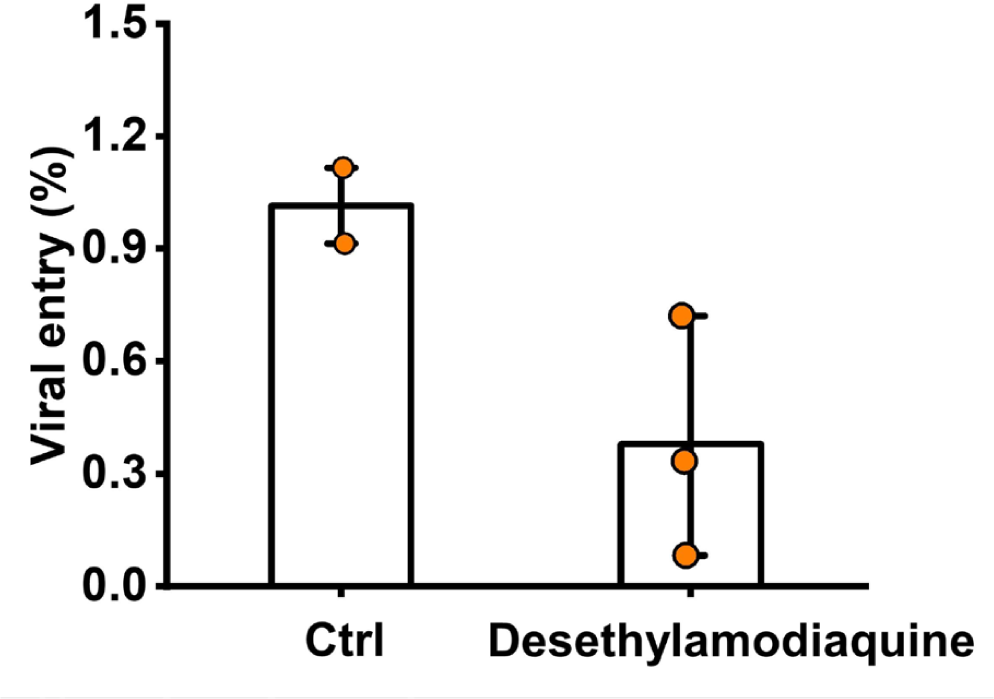
Effects of desethylamodiaquine on pseudotyped SARS-CoV-2 viral entry in human Airway Chips. Desethylamodiaquine was delivered into apical and basal channels of the chip at its C_max_ in human blood (1 μM), and one day later chips were infected with SARS-CoV-2pp while in the continued presence of the drugs for 2 days. The epithelium from the chips were collected for detection of viral gene by qRT-PCR; viral entry in untreated chips was set as 100%.

**Extended Data Fig. 9.**
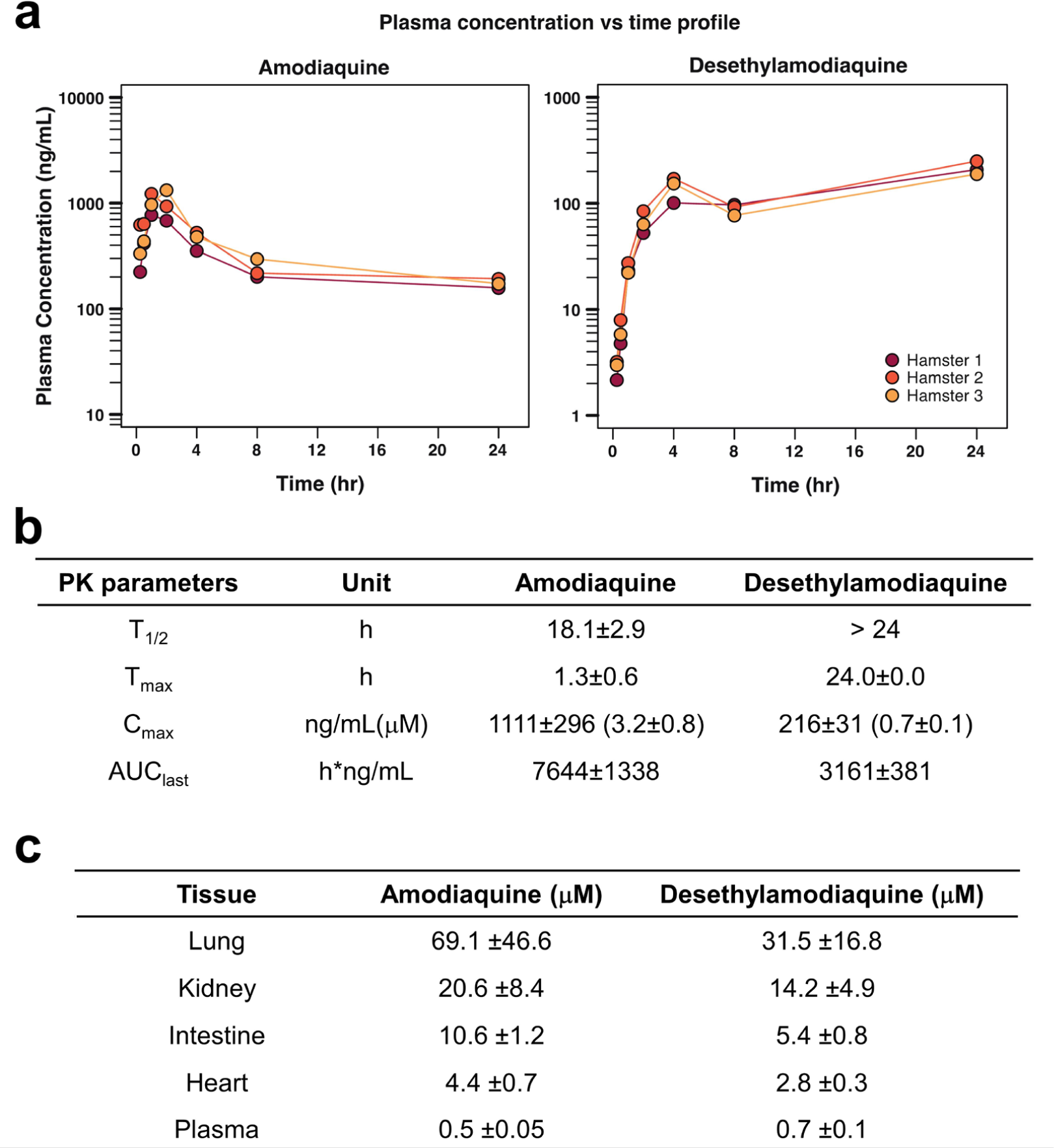
PK profiles for amodiaquine and desethylamodiaquine in hamsters. (**a**) Plasma concentration-time profiles showing mean concentration (± s.d.) of amodiaquine (left) and desethylamodiaquine (right) at different time points after a single subcutaneous injection of amodiaquine (50 mg/kg). (**b**) PK parameters for amodiaquine and desethylamodiaquine in plasma based on results shown in (**a**). (**c**) Concentration of amodiaquine and desethylamodiaquine in tissues (lung, kidney, intestine, heart) and plasma 24 hours after subcutaneous dosing of 50 mg/kg amodiaquine.

## Supplementary Materials

**Supplementary Table 1.** Limitations of current *in vitro* viral infection models^10–12, 24^.

| MODELS | LIMITATIONS |
| --- | --- |
| **Cell lines**<br>**(A549, MDCK)** | Minimal viral replication without the addition of exogenous proteases |
|  | Cannot be used for analysis of virus tropism |
|  | Lack host immune, tissue-level, or organ-level responses |
|  | Do not mimic the *in vivo* phenotype of human lung cells and tissues |
| **Ex vivo culture of**<br>**human lung tissue** | Short viability (4-10 days) |
|  | Limited availability of resources and expensive |
|  | Uncontrolled region-to-region and donor-to-donor variation |
|  | Poor reproducibility of experimental results |
|  | Difficult to analyze mechanism of infection or host responses |
|  | Not possible to study viral evolution |
| **Human organoids** | Lack of physiologically relevant organ-level microenvironment |
|  | Difficult to access apical surface of the epithelium |
|  | Lack of air-liquid interface |
|  | Cannot study mucociliary clearance |
|  | Lacks endothelium and circulating immune cells |
|  | Absence of relevant mechanical cues (air flow, vascular flow) |
|  | Thick ECM gel complicates permeability and drug studies |

**Supplementary Table 2.** Summary of antibodies used in this study.

| Protein/Structure/Cell | Antibody | Vendor and Catalog |
| --- | --- | --- |
| Tight junction | Alexa Fluor 594 anti-ZO-1 | Life Technologies, Cat# 339194 |
| Cilia | Alexa Fluor 647 anti-Acetyl-α-Tubulin | Cell Signaling Technology, Cat#81502 |
| VE-Cadherin | FITC anti-human VE-cadherin | BD Biosciences, Cat# 560411 |
| Goblet cell | Anti-Mucin5AC | Santa Cruz Biotechnology, Cat# sc-21701 |
| Club cell | Anti-human Uteroglobin/cc-10 | R&D Systems, Cat# MAB4218SP |
| Basal cell | Anti-Cytokeratin 5 | Sigma-Aldrich, Cat# SAB5300265 |
| Influenza NP | Anti-influenza NP | Invitrogen, Cat# MA516291 |
| hACE2 | Anti-hACE2 antibody | Abcam, cat. #ab239924 |
| ECM/Collagen | Anti-Collagen IV α1 | Novus Biologicals, Cat# NBP1-97716G |
| Neutrophil | Alexa Fluor 594 anti-human CD45 | Biolegend, Cat# 368520 |
| Influenza HA1 (H1N1) | Rabbit anti-influenza A H1N1 HA1 antibody | Sino Biological, Cat# 11692-T62 |
| Influenza HA (H3N2) | Mouse monoclonal [AT1B7] to Influenza A H3N2 HA antibody | Abcam, Cat# ab139361 |
| TMPRSS2 | Mouse anti-TMPRSS2 antibody | Novus Biologicals, Cat# H00007113-B01P |
| TMPRSS4 | Rabbit anti-TMPRSS4 antibody | Novus Biologicals, Cat# NBP1-56991 |
| TMPRSS11D | Mouse anti-TMPRSS11D antibody | Abnova, Cat# H00009407-B01 |
| TMPRSS11E | Rabbit anti-TMPRSS11E (DESC1) antibody | OriGene Technologies, Cat# TA350522 |
| Secondary antibody | Goat anti-mouse IgG, Alexa Fluor 488/594/647 | Life Technologies |
|  | Goat anti-rabbit IgG, Alexa Fluor 488/594/647 | Life Technologies |
|  | Goat anti-rabbit IgG H&L (HRP) | Abcam |
|  | Goat anti-mouse IgG H&L (HRP) | Abcam |

**Supplementary Table 3.**
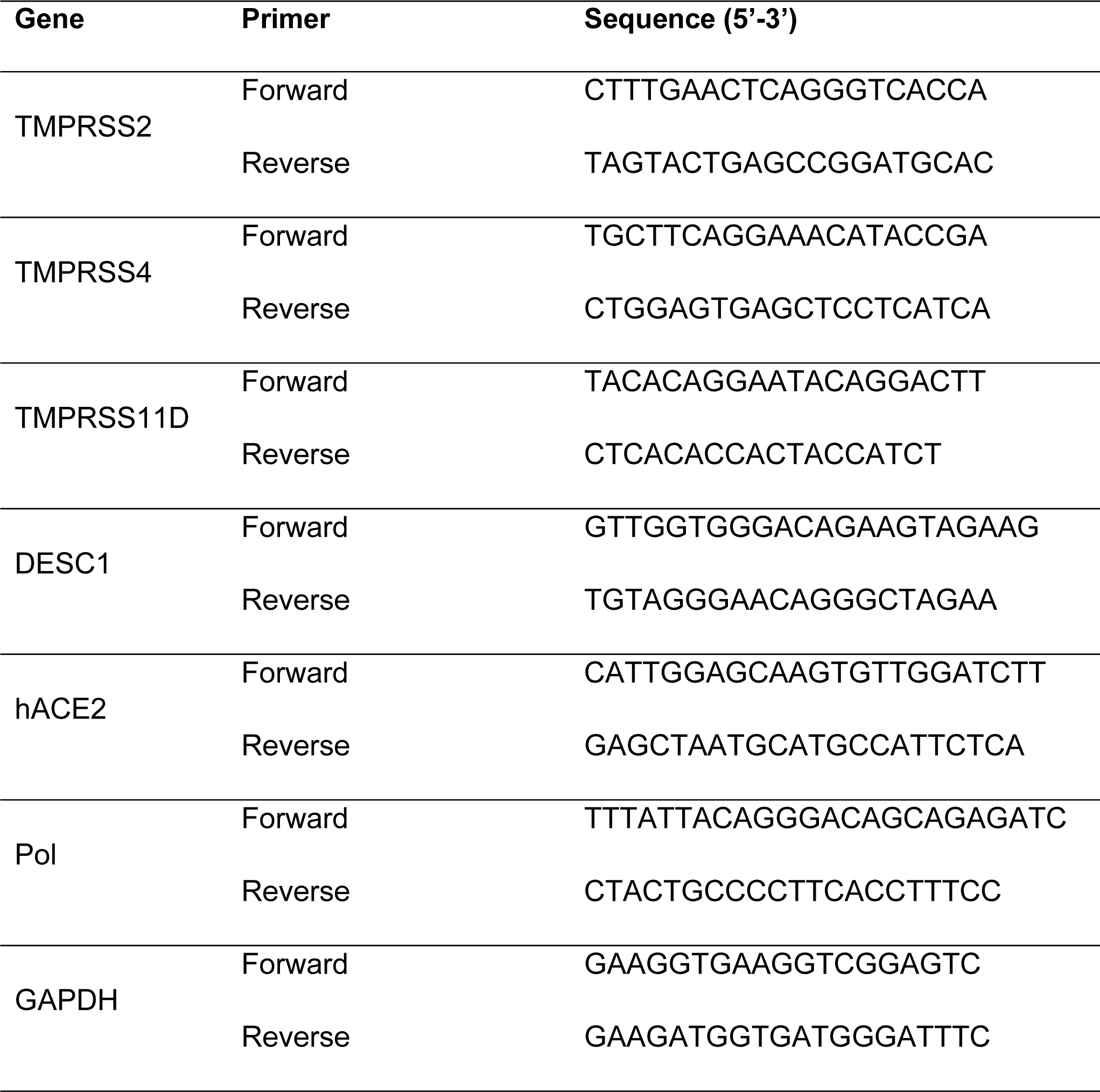
Primer sequences used for RT-qPCR analysis in this study.

### Other Supplementary Materials for this manuscript

**Movie S1 (.mp4 format). Time-lapse video recording showing infection of the human Airway Chip by GFP-labeled influenza PR8 (H1N1) virus recorded over 36 hours.** The human Airway Chip was inoculated with GFP-labeled PR8 virus (MOI = 0.01) and cultured for 36 h (images were recorded every 15 min). Virus infection is indicated by the progressive increase in GFP-positive cells (movies are played at 18,000 times real time). (**a**) GFP signal, (**b**) merge of GFP and bright field.

**Movie S2 (.mp4 format). Real-time imaging showing recruitment of human neutrophils to endothelium under flow in the human Airway Chip infected with influenza virus.** (**a**) The movie shows fluorescently-labelled human neutrophils flowing over a quiescent endothelium within the control Airway Chip that contains an airway epithelium on the opposite side of the porous membrane from the endothelium. Note that the neutrophils flow by and do not stick to the inactivated endothelium under these control conditions, as observed in normal vessels *in vivo*. (**b**) In contrast, many of the flowing neutrophils adhere to the surface of the activated endothelium within an Airway Chip that has been infected with influenza H1N1 virus (MOI = 0.1) via its introduction into the upper air channel, much as they do at sites of inflammation *in vivo*. The movies are played at 25 times real time.

## Notes

### Competing Interest Statement

D.E.I. is a founder and holds equity in Emulate Inc., and chairs its advisory board. D.E.I., L. S., R. P., K. H. B., H. B., and M. R. are inventors on relevant patent applications submitted by Harvard University.

### Summary of Updates

We further confirmed the drug efficacy against native SARS-CoV-2 in animal models.

